# Active Site-Directed Probes for targeting Bacterial Phosphoarginine Phosphatases

**DOI:** 10.64898/2026.01.13.697720

**Authors:** Christian E. Stieger, Yassmine El Harraoui, Luke Brennan, Michael Lisurek, Jan Vincent V. Arafiles, Charlotte Völkel, Kristin Kemnitz-Hassanin, Han Sun, Stephan Sieber, Christian P.R. Hackenberger

## Abstract

Canonical protein phosphorylation patterns are a thoroughly studied post-translational modification (PTM) driving distinct regulatory mechanisms in both prokaryotes, and eukaryotes. In contrast, the identification and investigation of essential components that regulate non-canonical phosphorylation has received considerably less attention, although these PTMs are associated with important functions. One notable example is arginine phosphorylation which modulates processes such as protein degradation, transcriptional regulation and spore germination in bacteria. Herein we introduce the first in class covalent activity-based probes to study phosphoarginine-phosphatases. We identify unsaturated phosphonamidic acids as bespoke electrophilic phosphoarginine (pArg) mimics, which allowed to uncover a series of unprecedented pArg-phosphatases, which in part had been previously annotated as low molecular weight tyrosine-phosphatases across phylogenetically distinct microbial species. This work, which serves as the first example of proteome-wide activity-based profiling of pArg phosphatases will help inform the development of new therapeutic modalities and expand our understanding of bacterial signal transduction.

## Introduction

Protein phosphorylation on serine, threonine, and tyrosine residues is well-known as a dynamic regulatory mechanism in both eukaryotic and prokaryotic organisms.^1–3^ In contrast, arginine phosphorylation (pArg), characterized by a chemically labile phosphoramidate bond, represents a distinct class of modification both chemically and biologically.^4,5^ While free phosphoarginine is a well-established phosphagen for energy storage in invertebrates, the post translational phosphorylation of arginine side chains within proteins has various biological functions. Following the discovery of this unique post-translational modification (PTM) in Gram-positive bacteria, pArg was soon linked to stress responses, in transcriptional control and protein quality control systems.^6–8^ In *Bacillus subtilis*, the kinase McsB selectively phosphorylates arginine residues (**Figure 1a**) on misfolded or damaged proteins, tagging them for degradation by the ClpC/ClpP protease complex.^9,10^ This process is tightly governed by the phosphatase YwlE, which reverses pArg modifications and maintains protein homeostasis (**Fig. 1a**).^11^ Knockout studies in *B. subtilis* show widespread accumulation of pArg in a Δ*ywlE* mutant, but not in wild-type cells highlighting YwlE’s central role in dephosphorylation.^6^ Additionally, pArg dephosphorylation by YwlE was discovered to be an essential step in spore germination.^12^ Structurally, YwlE is classified within the low-molecular-weight protein tyrosine phosphatase (LMW-PTP) family, which features a set of conserved active site residues, including a catalytic cysteine.^13^ The hydrolysis of phospho-arginine is facilitated by the recognition of its guanidinium group by Asp118 and Thr11 which positions the phosphate group in an ideal position for nucleophilic attack by Cys7. The phospho-cysteine intermediate is ultimately hydrolyzed to complete the catalytic cycle (**Fig. 1b**).^14^

**Figure 1:**
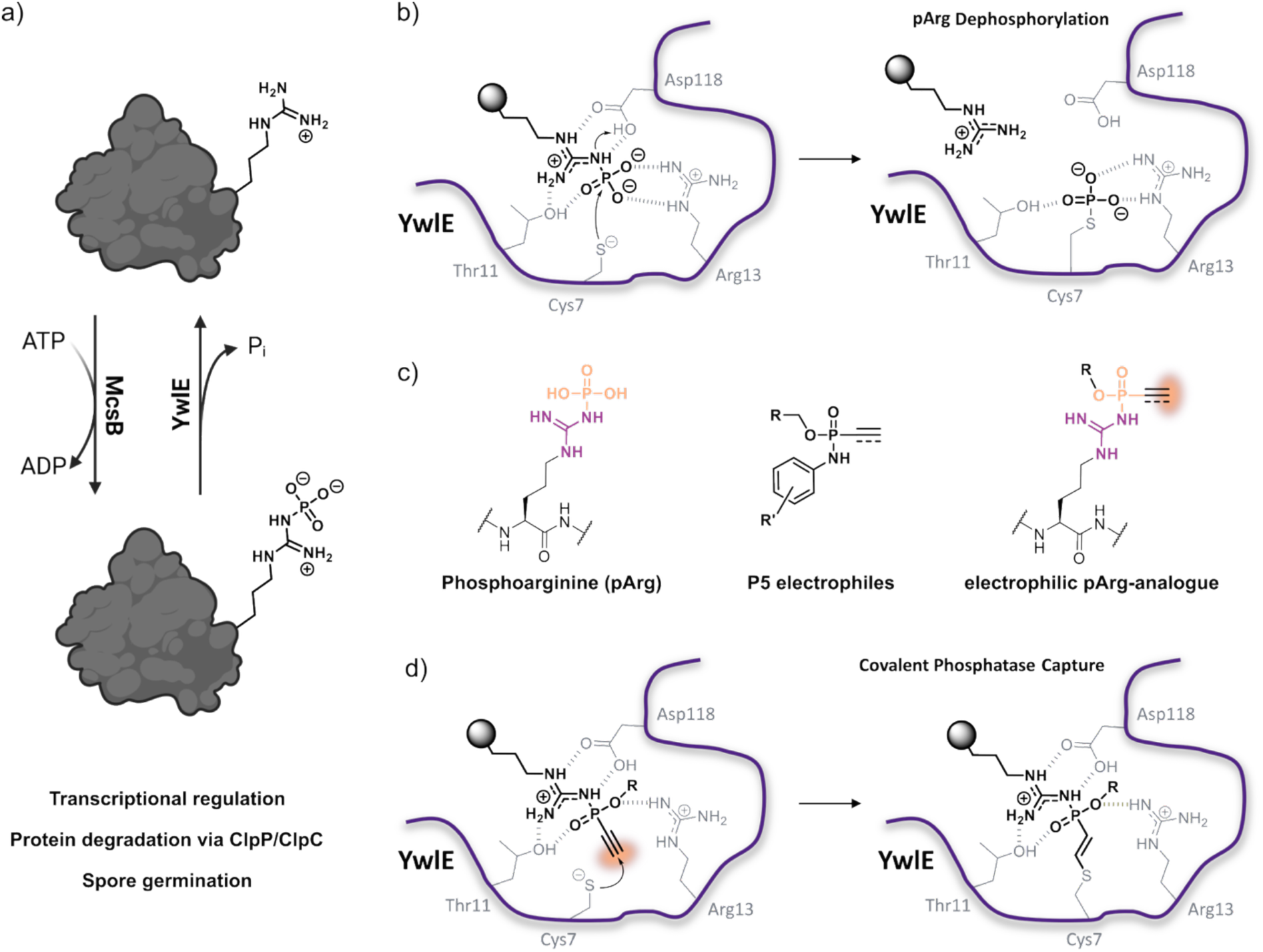
(a) Dynamic regulation of arginine phosphorylation and dephosphorylation by the corresponding kinase (McsB) and phosphatase (YwlE). **(b)** Schematic representation of the supposed reaction mechanism of YwlE, highlighting the active site residues involved in substrate recognition. **(c)** Chemical structure of phosphoarginine (left), ethynyl-phosphonamidate electrophiles for cysteine labelling (**P5**, middle) and proposed structure of electrophilic pArg mimetic (right). **(d)** Proposed mechanism of covalent capture of the active site cysteine (Cys7) of YwlE by electrophilic probes.

Available tools to study pArg-phosphatases have been restricted to photoaffinity-labelling probes derived from electrostatic pArg-mimetics^15^, pArg-based fluorescent activity probes^16^ and redox based inhibitors.^17^ However, no covalent activity-based probes targeting the active site of this unique enzyme class have been reported so far, limiting direct profiling of enzymatic activity. Indeed, covalent activity-based probes are useful for their ability to permit direct profiling of active, functional enzyme variants and not just binding motifs – highlighting their critical role as tools in functional proteomics. In contrast, there are several reports on activity-based probes for mechanistically related protein-tyrosine-phosphatases (PTPs).^18–21^ However, the chemistries used in these probes are inherently instable in neutral buffers^18–22^, show significant off-target reactivity^18–22^ or require a peptide motif for successful target recognition.^23,24^ Our group has previously reported on the use of unsaturated P^V^-electrophiles as cysteine-selective electrophiles for peptide- and protein-bioconjugation.^25–27^ Moreover, we could demonstrate that less reactive unsaturated P^V^-electrophiles can be incorporated into drug-derived ligands resulting in target selective covalent inhibitors.^28^ Given the shared geometry of the P^V^-center in pArg and phosphonamidate electrophiles, we reasoned that phosphonamidate-based pArg-analogues could be used as active site directed probes for this enzyme class (**Fig. 1c** & **d**). After structural optimization we discovered a core phosphonamidate motif suitable for chemical probe development. Several functional probes were able to selectively label recombinant YwlE from *G. stearothermophilus.* Moreover, the developed probes showed enrichment and identification of a series of pArg phosphatases from different Gram-positive bacteria. Among the most enriched proteins we also identified pTyr phosphatases, like *S. aureus* PtpA, and verified that these enzymes are putative pArg phosphatases using synthetic peptide substrates. In conclusion, we introduce a unique chemical entity for the selective detection and functional characterization of pArg phosphatases.

## Results

To enable the chemical interrogation of bacterial arginine phosphatases, we set out to design an active-site–directed electrophilic probe. Given the structural similarity between the native phosphoramidate moiety of phosphoarginine and unsaturated phosphonamidates, we hypothesized that vinyl- or ethynyl-phosphonamidate ethyl esters could serve as effective covalent probes by targeting the conserved catalytic cysteine found in known pArg phosphatases like YwlE_G.st._. To test this hypothesis, we started with the development of a reliable synthetic route towards guanidinium based unsaturated phosphonamidite electrophiles. For this purpose, we used the readily available ethyl vinylphosphonochloridate and guanidinium model compound **1** to identify suitable coupling conditions. Initial attempts using commonly employed organic bases, including triethylamine (TEA) and diisopropyl-ethylamine (DIPEA), failed to promote the coupling reaction. In contrast, treatment with 1,8-diazabicyclo[5.4.0]undec-7-ene (DBU) led to rapid consumption of the starting materials. However, alongside the desired phosphonamidate product, we observed the formation of the DBU adduct of **P1a** as a significant side product (Scheme S1). In quest for an alternative, we tested 2-tert-butyl tetramethyl guanidine, also referred to as Barton’s base, that has previously been reported as a strong, non-nucleophilic base for the functionalization of arginine containing peptides and natural products with NHS-esters.^29,30^ Gratifyingly, the use of Barton’s base resulted in full conversion of the starting material to the desired product and allowed isolation of **P1a** in 71% yield. Using the same conditions, we also accessed the alkyne version (**P1b,** see supplementary discussion on the synthesis of **P1b** and scheme S3). With these molecules in hand, we wanted to test their reactivity towards the previously reported and well characterized pArg-phosphatase YwIE from *G. stearothermophilus* (YwIE_G.st._), which was obtained by recombinant expression. Unfortunately, neither of the two compounds showed a detectable amount of labelling after overnight incubation under physiological conditions (PBS, pH 7.4, room temperature, 20 eq. **P1a** or **P1b, Fig. 2a**). Also, prolonged incubation times and elevated pH did not result in the formation of the desired adduct. To gain a better understanding of the interaction between our proposed probe and YwIE_G.st._, we performed molecular docking. Interestingly, we could not obtain a docking pose with the probe inside the active site pocket. Additionally, covalent docking between Cys7 and the terminal alkyne carbon resulted in several steric clashes between the probe and amino acid side chains (**Extended Data Fig. 1**). Altogether, the obtained results suggest that the additional steric demand of the phosphorus ethyl substituent compared to the canonical phosphate group is too big to be accommodated in the active site of the enzyme (**Fig. 2b** & **Extended Data Fig. 1**). Based on the literature, the combination of the positively charged guanidinium group as well as a negatively charged head group is essential for the recognition of the zwitterionic pArg by the enzyme.^11^ Therefore, we thought to generate the corresponding guanidino-phosphonamidic acid variants **P2a** & **P2b**. According to the docking studies, these groups would fit far better into the active site, boosting the likelihood of covalently engaging the active site cysteine (**Extended Data Fig. 2**).

**Figure 2:**
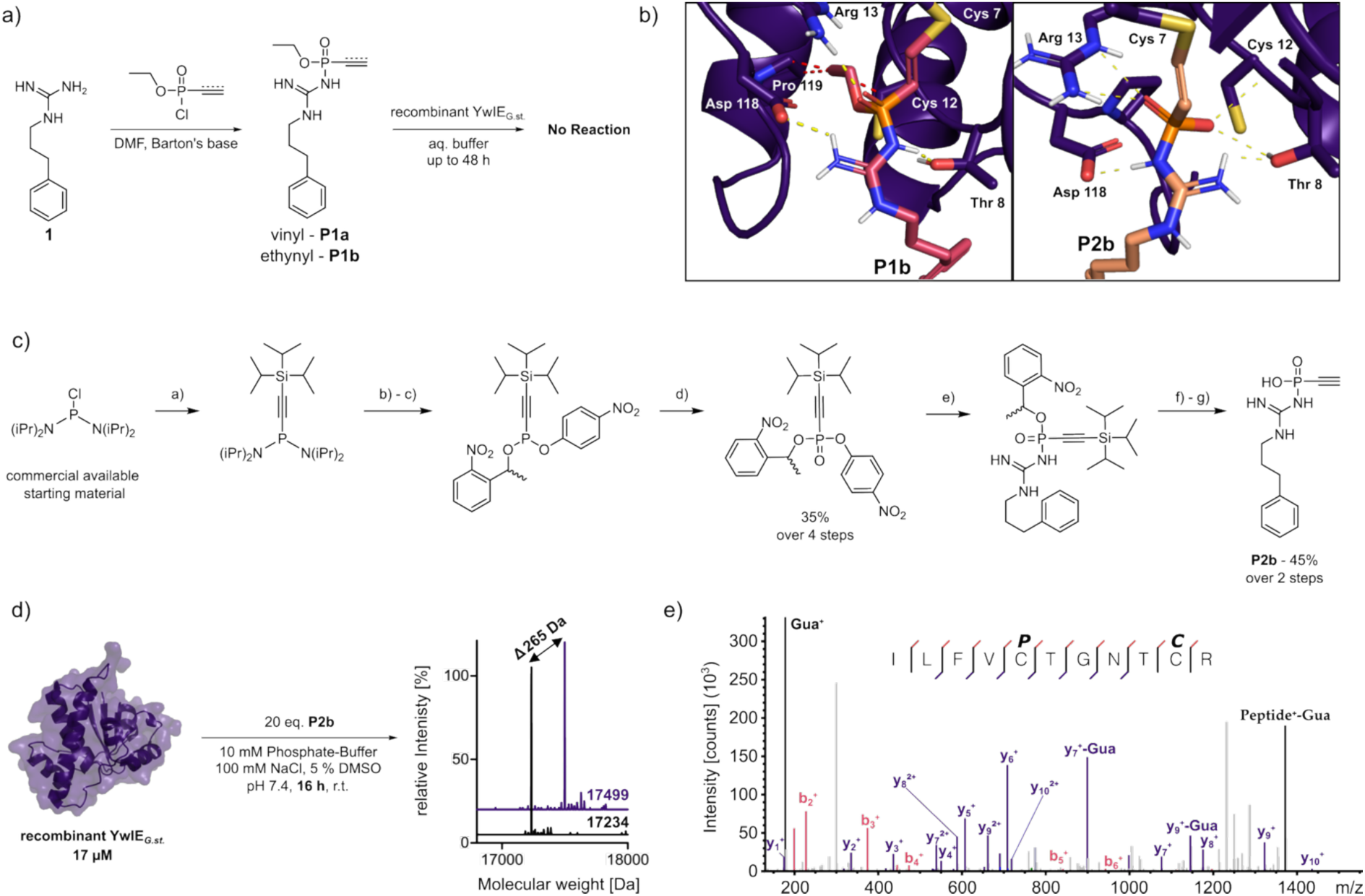
(a) Synthetic route towards compounds **P1a** and **P1b** and schematic representation of their reaction with YwlE. **(b)** Covalent docking structures of **P1b** and **P2b** onto a crystal structure of B. subtilis YwlE (PDB: 4KK3). Yellow lines indicate putative hydrogen bonds, red lined indicate steric clashes. **(c)** Synthetic route towards pArg-phosphatase probe **P2b**. a) 1.1 eq. TIPS-acetylene, 1.1 eq. BuLi, THF, -78 °C – r.t. b) 1.1 eq. tetrazole in MeCN, 1 eq. 1-(2-nitrophenyl)ethan-1-ol, -78 °C – r.t. c) 1.1 eq. tetrazole in MeCN, 1 eq. p-nitrophenol, -78 °C – r.t. over night. d) 3 eq. Luperox. e) 2 eq. Barton’s base, 1 eq. **1**, DMF, 5 min r.t. f) hv in MeCN, 30 min. g) 10 eq. NH_4_F in DMSO. **(d)** Schematic representation of the reaction between **P2b** and YwlE_G.sp._ and deconvoluted intact mass spectra of the unlabelled/labelled protein after 16 h. **(e)** Tandem MS/MS spectrum identifying the catalytic Cys7 as the site of labelling.

The vinyl-functionalized probe **P2a** could be accessed from bis(2-nitrobenzyl) vinylphosphonate in a straightforward manner. Selective mono-hydrolysis using aqueous NaOH followed by chlorination afforded the corresponding vinylphosphonochloridate, which reacted cleanly with guanidine model compound **1** in the presence of Barton’s base. Subsequent UV-mediated deprotection furnished the desired guanidinium-based vinylphosphonamidate probe **P2a** (Scheme S5). In contrast to the vinyl analogue, the synthesis of the ethynylphosphonamidate variant presented several challenges. Initial attempts to perform basic hydrolysis of bis(2-nitrobenzyl) ethynylphosphonate led to low yields and poor product purity. As an alternative, (2-nitrobenzyl) ethynylphosphonic acid (**6b**) was successfully accessed via acidic hydrolysis of a protected phosphonamidate precursor (Scheme S6). However, activation of **6b** using oxalyl chloride proved unreliable and inconsistent, often resulting in incomplete conversion or decomposition. To overcome this bottleneck, we envisioned the development of a bench-stable, fully functionalized P^V^ electrophile that could be stored and handled under ambient conditions and react directly with guanidine-containing substrates. While P^V^-chlorides are commonly used as activating groups, their high reactivity often limits stability. P^V^**-**fluorides have been reported as more stable alternatives, yet their use can be limited by toxicity and handling concerns. To address these issues, we turned to a strategy reported by Dutra et al., who described the use of para-nitrophenyl (pNP) phosphonates as air-stable precursors for the generation of fluorophosphonates.^31^ Given the strong electron-withdrawing character of the p-nitrophenyl group, we reasoned that a pNP-substituted ethynyl-phosphonate could serve as a viable and bench-stable intermediate. However, to prevent nucleophilic cross-reactivity of the para-nitro-phenolate leaving group, we decided to incorporate an additional TIPS-protecting group on the alkyne. Indeed, the corresponding pNP reagent was synthesized in a convenient one-pot, four-step sequence in 35% overall yield (**Fig. 2c**). By employing this reagent in the Barton’s base mediated reaction with **1**, we did observe full conversion to the desired product after only 5 minutes. Following UV-deprotection, NH_4_F treatment gave the desired probe **P2b** in 45% yield over two steps.

Overnight incubation of YwlE_G.st._ with the two phosphonamidic acid compounds resulted in a mass-shift of 265 Da indicative for covalent labelling with compound **P2b**, while no labelling could be observed for the vinyl variant **P2a** (**Fig. 2d** & **Extended Data Fig. 3**). To confirm the site of covalent modification, we subjected the labelled phosphatase to bottom-up LC-MS/MS proteomic analysis. Peptide mapping localized the modification at both active-site cysteine residues, with a slight preference for Cys7 over Cys12, in line with the catalytic mechanism proposed for YwlE_G.st._ (**Fig. 2e** & **Extended Data Fig. 4**). To further assess its selectivity and reactivity, we incubated **P2b** with a cysteine bearing EGFP mutant and TEV-protease containing an active site cysteine and observed no reaction. Additionally, stability studies in buffer revealed that compound **P2b** remained intact and reactive over relevant time scales, further supporting its suitability as a covalent probe.

Following this initial proof-of-concept, we sought to develop phosphonamidate probes equipped with functional handles that would enable downstream analysis via gel- and mass spectrometry-based activity-based protein profiling (ABPP). Starting from *N*-Boc-hexanediamine, we synthesized probes containing biotin (**P3**), fluorophores (e.g. fluorescein **P4**, TAMRA **P5**) or a bioorthogonal tetrazine functionality (**P6**, **Fig. 3a**, Scheme S9). All probes showed significant labelling of recombinant YwIE after overnight incubation. However, depending on the functional group stark differences in labelling efficiency were observed. While biotin- and fluorescein-functionalized probes achieved quantitative labelling after overnight incubation, the TAMRA- and tetrazine-tagged variants showed moderate labelling efficiencies of 60% and 30%, respectively (**Fig. 3b** & **Extended Data Fig. 5**).

**Figure 3:**
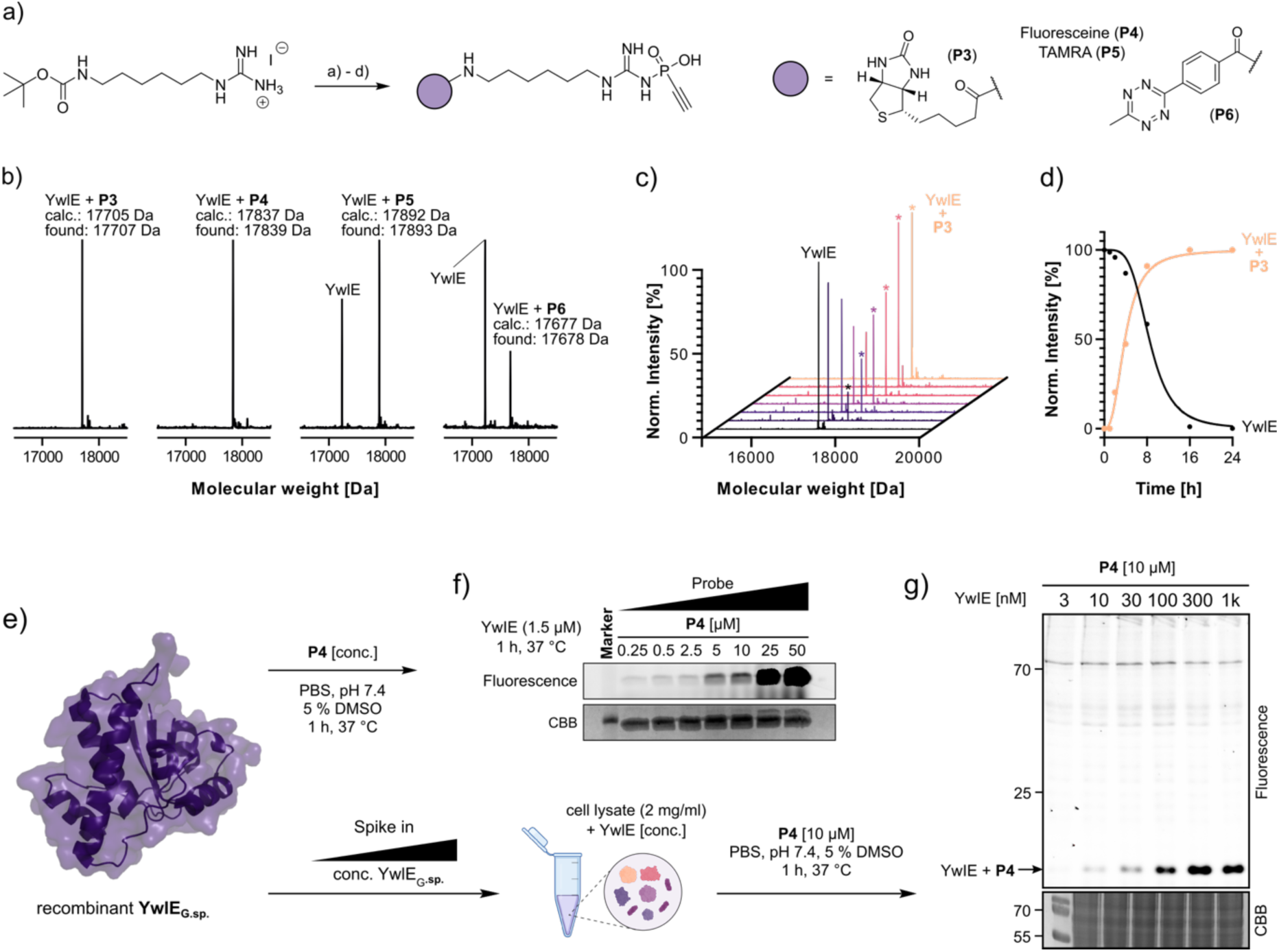
(a) Synthetic route towards functionalized pArg-phosphatase probes **P3**-**P6**. a) 2 eq. Barton’s base, 1 eq. **1**, DMF, 5 min r.t. b) TFA/DCM 1:1, 30 min, then 1 eq. HATU 3 eq. DIPEA and 1 eq. acid precursor c) hv in MeCN, 30 min. d) 10 eq. NH_4_F in DMSO. (for fluorophore containing probes, b) and c) were switched). **(b)** Deconvoluted intact MS spectra of YwlE_G.sp._ after 24 h incubation with the corresponding probe. **(c)** & **(d)** Labelling time-course of the reaction of YwlE_G.sp._ with probe **P3**. The peak corresponding to the labelled product is marked with an asterisk. **(e)** Crystal structure of YwlE_G.sp._ (PDB: 4KK3). Gel-based activity based protein profiling (ABPP) of recombinant YwlE_G.sp._ using increasing concentrations of **P4**. **(g)** Gel-based activity based protein profiling (ABPP) of increasing concentrations of recombinant YwlE_G.sp._ spiked into HEK293 cell-lysate using **P4**.

To gain deeper insight into probe reactivity and kinetics, we performed a time-course labelling experiment using only a minimal excess of fluorescent probe **P4** and biotin-probe **P3**. Intact protein mass spectrometry revealed a prominent mass shift corresponding to covalently labelled YwlE_G.st._ after only 1 hour (**Fig. 3c** & **Extended Data Fig. 6**). The labelling progressed in a time-dependent manner, reaching complete conversion after 16 hours (**Fig. 3c-d** & **Extended Data Fig. 6**). Given the robust labelling observed at early time points and under near-stoichiometric conditions, we next sought to define optimal conditions for gel-based ABPP with shorter incubation times. To this end recombinant YwlE_G.st._ (1.5 µM in PBS, pH 7.4) was incubated with decreasing concentrations of fluorescent probe **P4** for 1 h at 37 °C. A detectable fluorescent signal was observed at probe concentrations as low as 250 nM, while most intense labelling occurred >10 µM (**Fig. 3e-f**). Interestingly, the Coomassie staining revealed a faint, higher-migrating band above the expected molecular weight of YwlE_G.st._, likely corresponding to the covalently modified enzyme. While labelling efficiency is one key parameter, selectivity is just as important for the probe performance, especially in complex biological environments. To evaluate this, we spiked increasing concentrations of recombinant YwlE_G.st._ in a background of human cell lysate (2 mg/ml). Since mammalian cells are not expected to contain pArg processing enzymes, the only detectable signal should stem from the spiked-in YwlE_G.st_.^32^ The lysate containing YwlE_G.st_ was incubated with 20 µM **P4** for one hour at 37°C. In-gel fluorescence analysis revealed an intense, dose-dependent fluorescent band corresponding to YwlE_G.st._, with minimal detectable off-target labelling at the tested probe concentration (**Fig. 3d**). This result underscores the selectivity of our pArg-phosphatase probes, even in the presence of a complex proteomic background, and supports its utility for selective detection and enrichment of pArg phosphatases from biological samples.

Having determined that already short incubation times can lead to a significant amount of labelled protein with robust selectivity, we wanted to test the developed probes in a more native environment. As a first model system we selected the *E. coli* used for recombinant expression of YwlE_G.st._ (BL21(DE3) *E. coli*). Although an analogue of the pArg kinase McsB has been identified in *E. coli*, Gram-negative bacteria are generally considered to lack the enzymatic machinery required for pArg turnover.^32,33^ Consequently, no *E. coli* protein should interact with our probe. At first, we verified that YwlE_G.st._ is expressed at a reasonable level without IPTG induction. Quantitative bottom-up proteomics of the bacterial lysate revealed that YwlE_G.st._ is stably expressed at the same level as other enzymes with active site cysteines (**Extended Data Fig. 7a**). Incubation of the bacterial cell lysate with **P4** probe for one hour yielded an intense fluorescent band around 15 kDa, indicative for YwlE_G.st._(**Fig. 4b**). Interestingly, two additional bands were detected at a higher molecular weight. However, these bands were not detected in case of the TAMRA-probe **P5** (**Fig. 4c**). This indicates that the observed labelling could stem from a certain affinity for Fluorescein, or auto fluorescent properties. Yet, to unambiguously verify the selectivity of our probe, we performed a proteomics pull-down experiment using the biotinylated probe **P3** (**Fig. 4d**). In brief, the lysate was incubated with 20 µM **P3** for 1 h at r.t. followed by acetonitrile precipitation, streptavidin enrichment and tryptic digestion (see Methods section for detailed procedure). To our delight, the arginine phosphatase YwlE_G.st._ was more than 16-fold enriched compared to the DMSO-treated control, demonstrating the probe’s suitability for chemo-proteomic profiling of pArg phosphatases in bacterial samples. In addition to YwlE_G.st._, a single unexpected protein hit (YqjG) was significantly enriched across replicates. We reason that this off-target labelling arises from the presence of a highly reactive active-site cysteine in YqjG, which may readily undergo reaction with electrophilic groups, even those with comparatively modest reactivity.^34^

**Figure 4:**
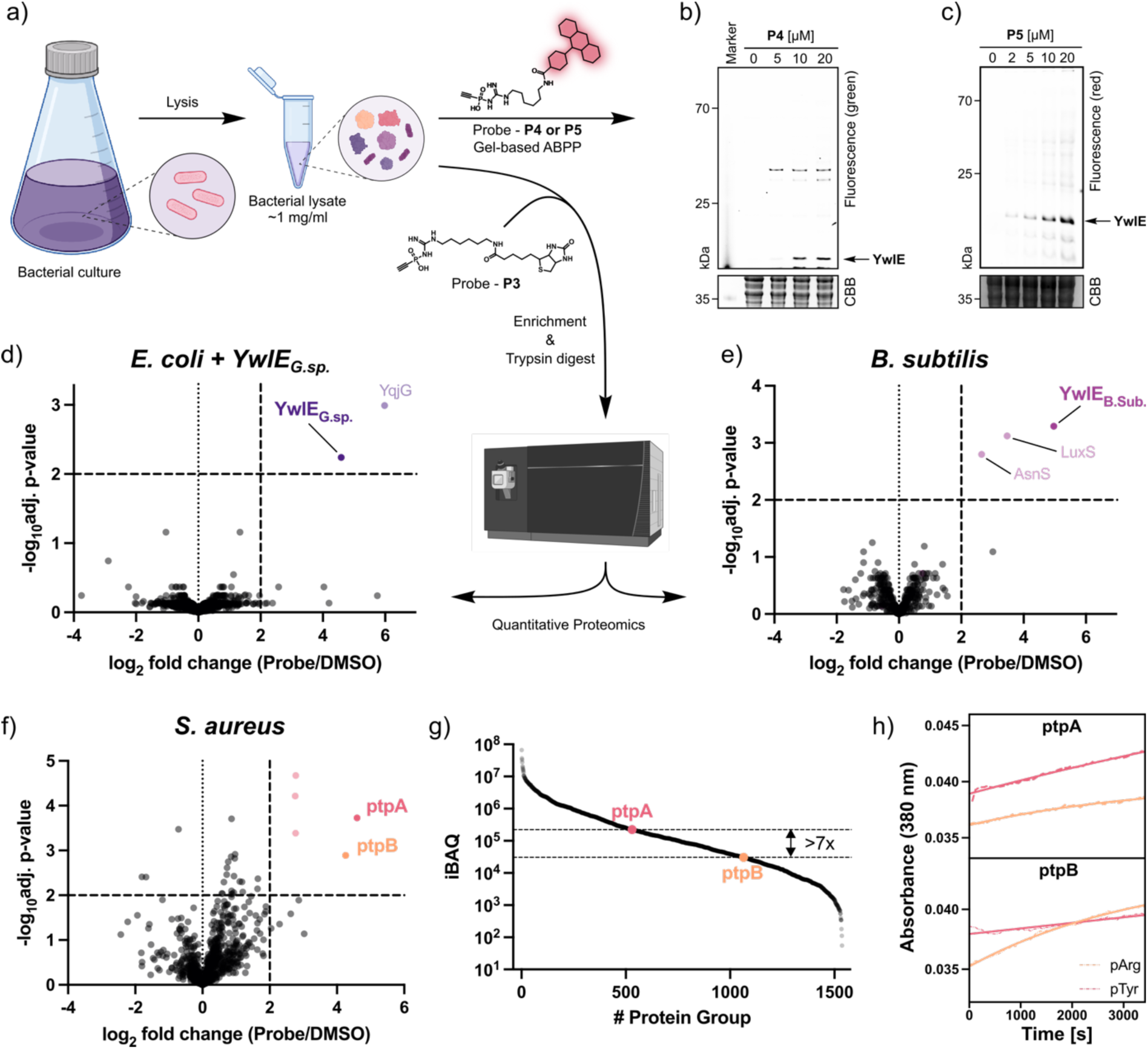
(a) Schematic representation of the preparation of bacterial lysate followed by incubation with functional probes **P3**-**P5**. **(b)** & **(c)** Gel-based activity based protein profiling (ABPP) in lysate from E. coli transfected with a plasmid for YwlE_G.sp._ using probe **P4** or **P5**, respectively. Pull-down proteomics from **(d)** E. coli transfected with a plasmid for YwlE_G.sp_ **(e)** B. subtilis and **(f)** S. aureus using biotinylated pArg-probe **P3**. Significance cut-off was set at adj. p-value < 0.01 and log_2_ fold change >2. Waterfall plot of all quantified proteins from S. aureus lysate based on iBAQ values. **(h)** Determination of the reaction rate for dephosphorylation of pTyr and pArg containing peptides by S. aureus PtpA and PtpB.

This remarkable selectivity prompted us to investigate the performance of our probe in non-genetically modified organisms. Arginine phosphorylation was first described as a widespread post-translational modification in *Bacillus subtilis*, which remains the most extensively studied organism with respect to this modification. In *B. subtilis*, pArg plays a central role in the cellular stress response, particularly under heat shock conditions, where misfolded proteins are tagged by arginine phosphorylation and subsequently targeted for degradation by the ClpC/ClpP protease complex.^6,9^ More recently, dephosphorylation of arginine residues has also been implicated in spore germination, pointing to additional regulatory functions beyond proteostasis.^12^ At first, we investigated the expression level of endogenous YwIE. Interestingly quantitative proteomics revealed that endogenous YwlE levels in the tested *B. subtilis* strain were significantly lower, compared to *E. coli* engineered to express YwlE_G.st._ (**Extended Data Fig. 7b**). Following incubation with biotin probe **P3** with subsequent streptavidin enrichment and on-bead trypsin digestion, we identified YwIE as the most significantly enriched protein with a fold_-_change >20 compared to the DMSO treated control (**Fig. 4e**). Notably, another protein that was highly enriched was the quorum sensing regulator LuxS. This finding may point to an unanticipated role of arginine phosphorylation in quorum sensing and intercellular communication in *B. subtilis*.^35,36^

Compared to *B. subtilis*, where all previous data suggests that YwIE is the sole pArg phosphatase, in other bacteria this situation is more ambiguous. In *Staphylococcus aureus*, for example, more than thousand unique arginine residues are reported to be phosphorylated, however, the landscape of pArg dephosphorylation is less clear.^32,37–39^ While serine/threonine phosphatase Stp1 was speculated to possess pArg-phosphatase activity recent studies reveled a more indirect role for this enzyme.^32^ To investigate this further, we applied our probe to native *S. aureus* lysate and performed chemoproteomic analysis. Surprisingly, not only the annotated pArg-phosphatase PtpB, but also its close homolog PtpA, previously described as a pTyr phosphatase, were significantly enriched (**Fig. 4f**). The role of PtpB in pArg dephosphorylation has been established both *in vitro* by Fuhrmann *et al.* and later *in vivo* by Junker *et al.* In contrast, PtpA has been reported to act primarily on phosphotyrosine residues, at least under *in vitro* conditions. To further dissect the functional relevance of this observation, we next assessed the global expression level of PtpA and PtpB in the tested *S. aureus* strain. Interestingly, we found PtpB to be almost 10-times less abundant than PtpA (**Fig. 4g**). While previous studies have indicated a substrate preference for either pArg or pTyr in the cases of PtpB and PtpA, respectively, residual cross-reactivity cannot be ruled out. To directly evaluate the substrate specificity of both phosphatases, we recombinantly expressed the two proteins and synthesized defined phosphopeptides containing either phosphoarginine or phosphotyrosine residues. Using these peptides, we performed kinetic dephosphorylation assays. As expected, PtpB exhibited strong activity towards the pArg-containing substrate, while PtpA showed a clear preference for pTyr. Notably, however, PtpA retained substantial catalytic activity towards the pArg substrate, suggesting a broader substrate scope than previously anticipated (**Fig. 4h**). Consistent with this observation, we also detected covalent labelling of recombinant PtpA with biotinylated probe **P3**, albeit to a lesser extent than seen for YwlE_G.st.._Based on these observations, we propose that substrate specificity alone does not fully account for the observed *in vivo* activity of PtpA and PtpB. Instead, the relative expression levels and local concentrations of both phosphatases and their substrates are likely to play a critical role in determining functional outcome. This suggests a context-dependent duality, where either enzyme may act on pArg or pTyr residues depending on the cellular environment. This might be an important consideration for future investigations in the pArg phosphoproteome. While previously, a ΔptpB strain enabled the first identification of pArg sites in *S. aureus*, the double deletion of PtpA and PtpB variant may provide a more comprehensive view of pArg in bacterial physiology.

Finally, we wanted to leverage our newly developed pArg-phosphatase probes to discover previously unannotated arginine phosphatases in other bacteria. *Listeria monocytogenes*, a Gram-positive pathogen associated with listeriosis, represents a phylogenetically distinct organism that is expected to harbor pArg-processing enzymes. Following the same enrichment strategy as described above, we profiled native *L. monocytogenes* lysate for the presence of potential pArg phosphatases. Remarkably, among the significantly enriched proteins, we identified lmo0938, an annotated tyrosine phosphatase whose active site closely resembles that of *S. aureus* PtpA (**Fig. 5a**-**b**). Moreover, another significantly enriched yet unannotated protein, lmo2540, displays high sequence similarity to *S. aureus* PtpB. These results suggest that *L. monocytogenes*, like *S. aureus*, may rely on multiple cysteine-based phosphatases to modulate its pArg signaling pathways.

**Figure 5:**
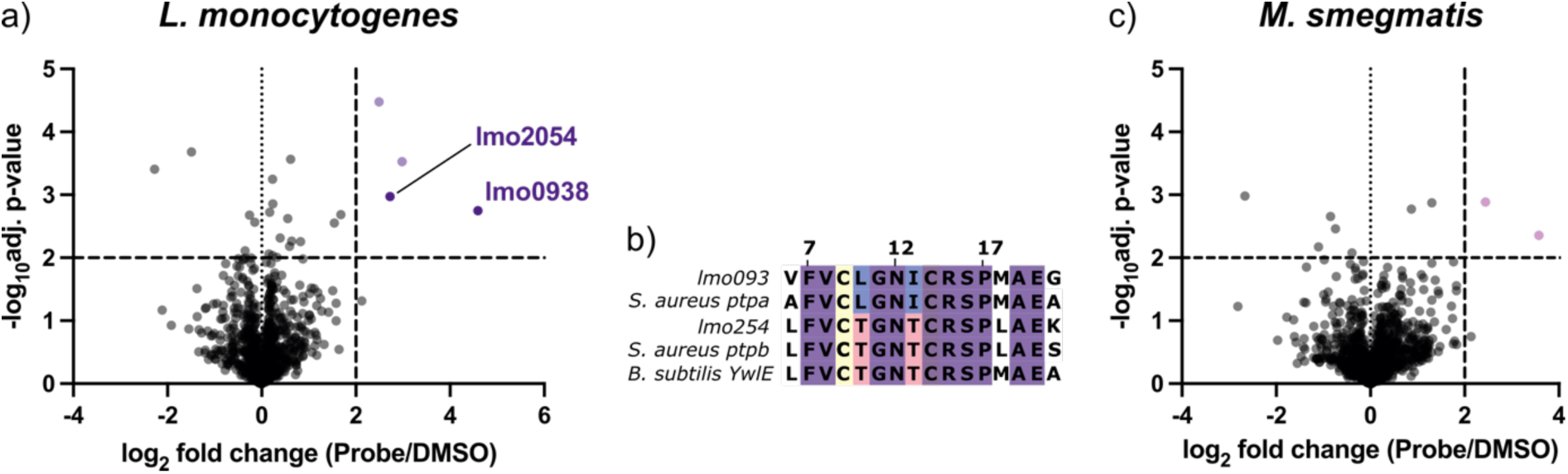
(a) Pull-down proteomics from L. monocytogenes lysate using biotinylated pArg-probe **P3**. **(b)** Sequence alignment of lmo0938 and lmo2054 with other reporter pArg/pTyr phosphatases. **(c)** Pull-down proteomics from L. monocytogenes lysate using biotinylated pArg-probe **P3**. Significance cut-off was set at adj. p-value < 0.01 and log_2_ fold change >2.

Beyond Gram-positive species, mycobacteria have also been proposed to utilize arginine phosphorylation. Indeed, a recent phosphoproteomics study identified several pArg sites in *Mycolibacterium smegmatis*, and a McsB-like arginine kinase has been annotated in the genome.^33^ However, until now no corresponding pArg-phosphatase has been identified. To address this gap, we applied our probe-based approach to native *M. smegmatis* lysates. In stark contrast to the findings in *B. subtilis*, *S. aureus*, and *L. monocytogenes*, no significantly enriched proteins were detected (**Fig. 5c**). This result supports previous speculation that mycobacteria may lack pArg phosphatase, at least such featuring an active-site cysteine, and suggests that alternative or non-canonical dephosphorylation mechanisms may be involved.

## Conclusion

In summary, we report the first development of a series of active-site directed covalent chemical probes designed to target and probe bacterial arginine phosphatases. Leveraging the highly modular, and exquisitely selective reactivity of unsaturated phosphonamidite warheads we could achieve efficient, selective labelling of recombinant pArg phosphatases such as YwlE_G.st._ even in complex environments, such as in human cell lysate. The fidelity of these probes was verified through a combination of molecular docking exercises, in-gel experiments, MS-based protein labelling experiments, and in chemo-proteomics experiments. Furthermore, we applied a biotinylated probe in the functional proteomic profiling of pArg phosphatases in a range of phylogenetically distinct microbial species (such as *B. subtilis, S. aureus,* and *L. monocytogenes*), implicating a role for a series of LMW-PTP’s in pArg de-phosphorylation processes. We reason these ABPP-probes will serve as critical functional tools to gain a deeper understanding of the role of pArg dephosphorylation in bacterial signal transduction and to investigate novel therapeutic approaches.

## Acknowledgment

We thank E. Poulou for valuable scientific discussion and B. Kindt, I. Kretzschmer and J. Lim for excellent technical assistance. We furthermore want to thank Dr. Peter Schmieder for assistance with NMR measurements. This work was supported by grants from the Deutsche Forschungs-gemeinschaft (DFG, SPP1623 HA 4468/10-1 and RTG GRK2473), the Leibniz Society (SAW-2018-FMP-4-P5label, T18/2017), the Einstein Foundation Berlin (Leibniz–Humboldt Professorship), the European Union (ERC, breakingBAC 101096911) and the TUM Innovation Network NextGenDrugs funded under the Excellence Strategy of the Federal Government and the Länder. C.E.S. was supported by a PhD fellowship of the Studienstiftung des Deutschen Volkes and J.V.V.A. was supported by the Alexander von Humboldt Research Fellowship for Postdocs. The mass spectrometry proteomics data have been deposited to the ProteomeXchange Consortium via the PRIDE^40^ partner repository with the dataset identifiers PXD072215 and PXD072717.

## Methods

### Chemical Synthesis

Details of the synthesis and characterization of all precursors, probes and peptides and supplementary discussion on the synthesis are given in Supplementary Information Section 2 & 3.

### Docking experiments

The binding mode of Ligands/Probes was modelled using the docking program GOLD (version 2023, Cambridge Crystallographic Data Center)^41^ integrated in the software package DiscoveryStudio (BIOVIA), the ChemPLP scoring function was employed.^42^ The protein structure 4KK3^11^ was taken from the PDB. The binding site was selected manually using the residues Cys7, Thr11, Arg13 and Asp118. For covalent restraints between the probe(s) and the protein Cys7 or Cys12 were selected. Docking poses were visualized using Pymol.

### Recombinant expression and purification of YwlEG.sp

BL21(DE3)-pLys cells were transformed with the corresponding plasmid that was obtained as a gift from Dr. Tim Clausen. A streak of colonies from a Luria Bertanni (LB)-agar plate was used to inoculate a 10 mL pre culture in Ampicillin containing LB medium and incubated overnight at 37°C under agitation. 1 L of Ampicillin containing LB medium was inoculated with the overnight culture and incubated at 37°C at 180 rpm until an OD_600_ ∼ 0.8 was reached. Cultures were then induced with 0.5 mM isopropyl ß-D-1-thiogalactopyranoside (IPTG) and further incubated for 3 h. Expression cultures were harvested by centrifugation (4000 xg, 30 min, 4°C) and resuspended in 15 ml phosphate buffer (50 mM, 300 mM NaCl, pH 7.4) supplemented with Benzonase, lysozyme and PMSF (2.5 mM). Bacteria were lysed by sonication (3x 5 min, 30% Amplitude) on ice followed by centrifugation to remove cell debris (25000 xg, 4 °C, 30 min). Crude lysate was loaded onto 5 mL HisTrap FF (GE Healthcare, USA) column via a BioRad NGC system (BioRad, USA), washed and eluted with increasing imidazole concentration (10-20-100 mM). Product containing fractions were combined and TEV-cleavage was performed overnight (1:10 w/w) in dialysis against PBS. Pure protein was obtained after size exclusion purification using a Superdex 75 10/300 GL column. YwlE containing fractions were analyzed by SDS-PAGE and high-resolution mass spectrometry (HRMS), combined, concentrated, aliquoted and stored at -70°C until further usage (Extended Data Figure 8).

#### Recombinant expression and purification of S. aureus PtpA/B

Bacterial expression plasmids (pET19b_PtpB and pETPhos_PtpA) were gifted by Prof. Virginie Molle. BL21(DE3) *E. coli* were transformed with the following plasmids by heat-shock and were plated onto LB-agar plates with carbenicillin. Single colonies were inoculated into 5-mL starter LB media cultures with carbenicillin and incubated overnight at 37°C with constant agitation. The starter cultures were added to 500 mL of LB medium with carbenicillin and were incubated at 37°C with constant agitation OD_600_ = 0.6. Expression was induced by adding IPTG to a final concentration of 0.5 mM and incubation at 16°C overnight. Induced cells were collected by centrifugation (4000 ×*g*, 15 min) and resuspended in 50 mM Tris-HCl, pH 7.4, 100 mM NaCl before lysis by sonication (3×, 2 min, 30% Amplitude). The lysates were clarified by centrifugation (25000 ×*g*, 30 min, 4°C) and purified using TALON^®^ metal affinity resins (Takara) with 50 mM Tris-HCl, pH 7.4, 100 mM NaCl, 200 mM imidazole. Fractions containing the proteins of interest were combined and were dialyzed and buffer exchanged to 20 mM HEPES pH 7.5, 150 mM NaCl, 10% glycerol. Purified proteins were concentrated by ultrafiltration (Amicon) to a final concentration of 1.5 mg/mL. Proteins were characterized by SDS-PAGE and HRMS, aliquoted and stored at -70°C until further usage (Extended Data Figure 9).

### Preparation of bacterial lysates for ABPP experiments

**E. coli** (BL21(DE3)-pLys) transfected with the plasmid for YwlE expression were grown in 5 ml LB-medium supplemented with Ampicillin over night at 37 °C with shaking at 200 rpm. The next day, 1 liter of medium was inoculated with two overnight cultures (total of 10 ml) and grown at 37 °C until an OD_600_ of 0.8 was reached. Subsequently, the bacteria were harvested by centrifugation (4000 g, 30 min, 4 °C) and the pellet was stored at -20 °C until further usage.

Right before lysis, the pellet was resuspended in 50 ml PBS (pH 7.4) and DNAse was added to the slurry. Bacteria were lysed by sonication (3x 2 min, 30% Amplitude) on ice, followed by debris centrifugation (25’000xg, 30 min, 4°C).

**B. subtilis** (DSM 347) were grown, harvested and lysed as described above, but TB-medium without antibiotics was used instead of LB-medium.

**HEK293** cells were cultured in 75 cm^2^ cell culture flasks to approx. 80% confluency. The cells were washed with PBS twice transferred to 1.5 ml Eppendorf-tubes and lysed by sonication using a Bioruptor (5 min, cycle 30/30 s, 4°C).

***S. aureus*** (USA300) cultures were initiated by inoculating 5 mL of B-medium with 5 µL of glycerol stock and incubated overnight at 37 °C with shaking at 200 rpm. The overnight cultures were then diluted 1:100 into fresh 50 mL B-medium and grown to the early stationary phase. Cells were harvested (6000 x g, 4 °C, 10 min) and the pellets were washed with PBS (10 mL, 6000 x g, 4 °C, 10 min). The Pellets were resuspended in 5 mL PBS, transferred to lysis tubes containing 0.1 mm ceramic beads (91-PCS-CK01L, *Peqlab*), and lysed in a Precellys 24 bead mill (3 x 30 s, 6500 rpm) while cooling with a liquid nitrogen cooled airflow and centrifuged (21100 x g, 1 h, r.t.).

***L. monocytogenes*** (EGD-e) were grown, harvested and lysed as described for *S. aureus*, but BHB-medium was used instead of B-medium.

***M. smegmatis*** (DSM43756) were grown, harvested and lysed as described for *S. aureus*, but 7H9 + OADC + 0.05% Tween-80 was used instead of B-medium and the overnight culture was grown for 48 h. Additionally 50 mM Tris/HCl was used as lysis buffer instead of PBS.

The concentration of the corresponding lysate was determined using a bicinchoninic acid (Pierce^TM^ BCA Protein Assay Kit, Thermo-Fisher Scientific) assay according to the manufacturer’s protocol. Lysates were either used immediately or stored at -70 °C.

### Lysate labelling for in gel ABPP

Lysates (approx. 100 μg of protein each) were incubated with the indicated concentration of fluorescent-pArg-phosphatase-probe **P4** or **P5** for 1 h at room temperature. Afterwards, proteins were precipitated with the 4-fold volume of acetonitrile and cetrifuged (14’00 rcf, 5 min). The solvent was removed and precipitates were resuspended in 80% EtOH followed by another round of centrifugation (10’000 rcf, 5 min). The pellets were aspirated and left to dry on air for 5-10 minutes. Afterwards, the precipitate was redissolved in 40 µl 8 M urea (100 mM Tris, pH 8), 8 µl Laemmli buffer were added and the samples were denatured at 95 °C for 5 minutes. Proteins were seperated via SDS-PAGE (12% or 4-20% acrylamide gels) and analyzed on a Gel Doc XR+ (Bio-Rad, USA). If indicated, lanes and bands were quantified in ImageLab 6.1.

### Lysate labelling sample preparation and data analysis for mass spectrometry based ABPP

Lysates were treated analogously to gel based ABPP experiments. Samples were prepared according to published procedures^43,44^ with minor adjustments. For detailed procedures, acquisition and analysis parameters see Supplementary Information Section 2.

### Phosphatase Activity Assay

The respective proteins investigated; PtpA, and PtpB were recombinantly expressed as described above. Phosphatase activity against phospho-peptides (for synthesis see Supplementary Information Section 3) was determined by the EnzCheck Phosphate assay (Invitrogen). In this assay, inorganic phosphate released by the phosphatase is a co-substrate/activator of the purine nucleoside phosphorylase that catalyzes a photometrically measurable reaction. The Reaction samples of a 100 μl working volume containing 20 mM Tris/HCl, pH 7.5, 50 mM NaCl, 200 μM 2-amino-6-mercapto-7-methylpurine riboside, 1 U/ml purine nucleoside phosphorylase, 100 μM of the phospho-peptide substrate and 1 μM phosphatase were pipetted in a 96-well plate. Reaction mixtures were pre-formed in the absence of the relevant phosphatase to account for background phosphate signal which may be present (5 min pre-incubation period), whereafter the reaction and monitoring thereof was initiated upon addition of the relevant phosphatase. The change in absorbance (380 nm) was monitored in 30s intervals, and apparent specific activities were directly plotted to visualise pArg vs pTyr specificity.

## Extended Data Figures

**Extended Data Fig 1:**
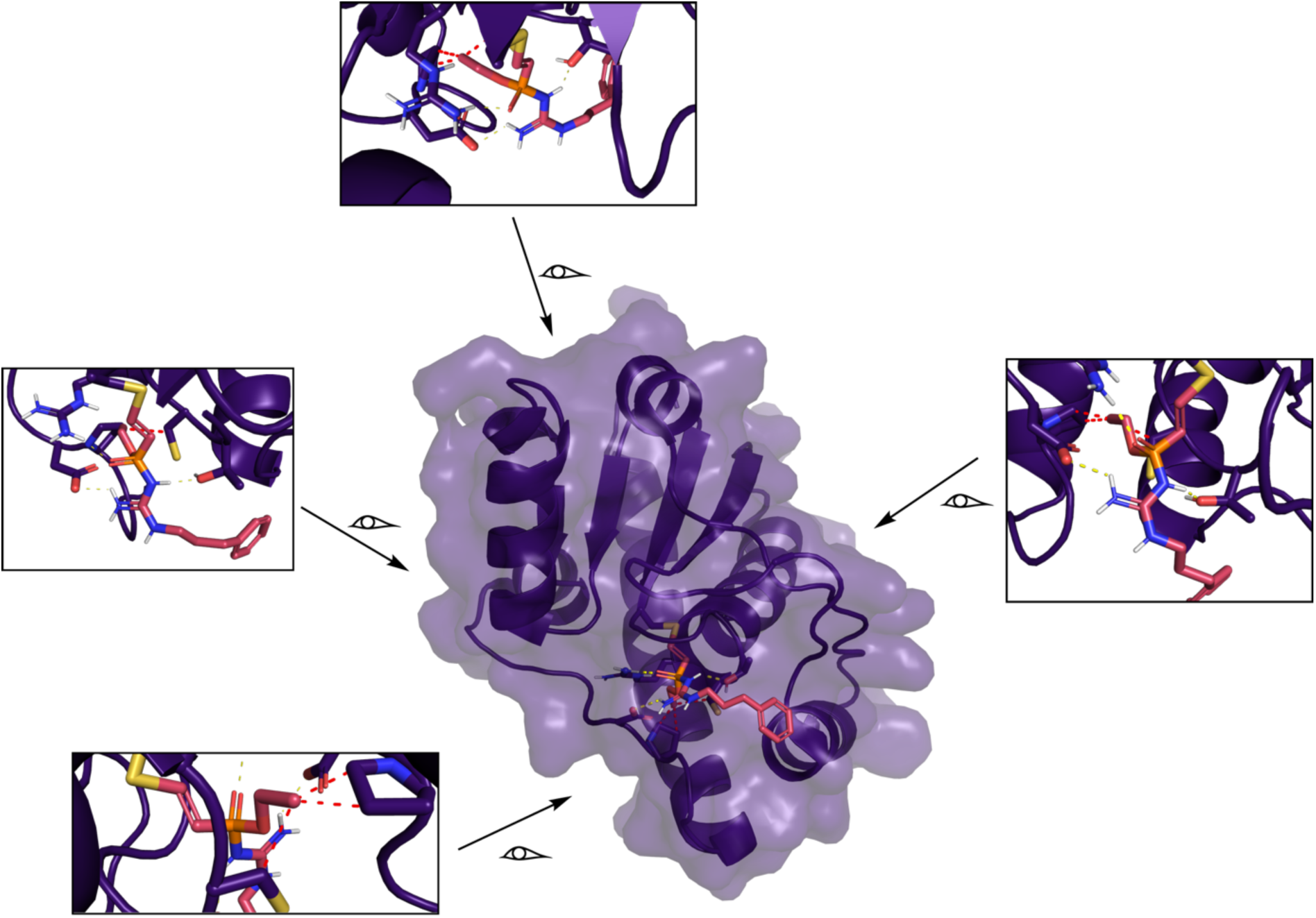
Covalent docking of P1b onto active site Cys7 of YwlE. (PDB: 4KK3). Yellow dotted lines indicate putative hydrogen-bonds. Red dotted lines indicate steric clashes between the O-ethyl substituent and Pro119 and Cys12.

**Extended Data Fig 2:**
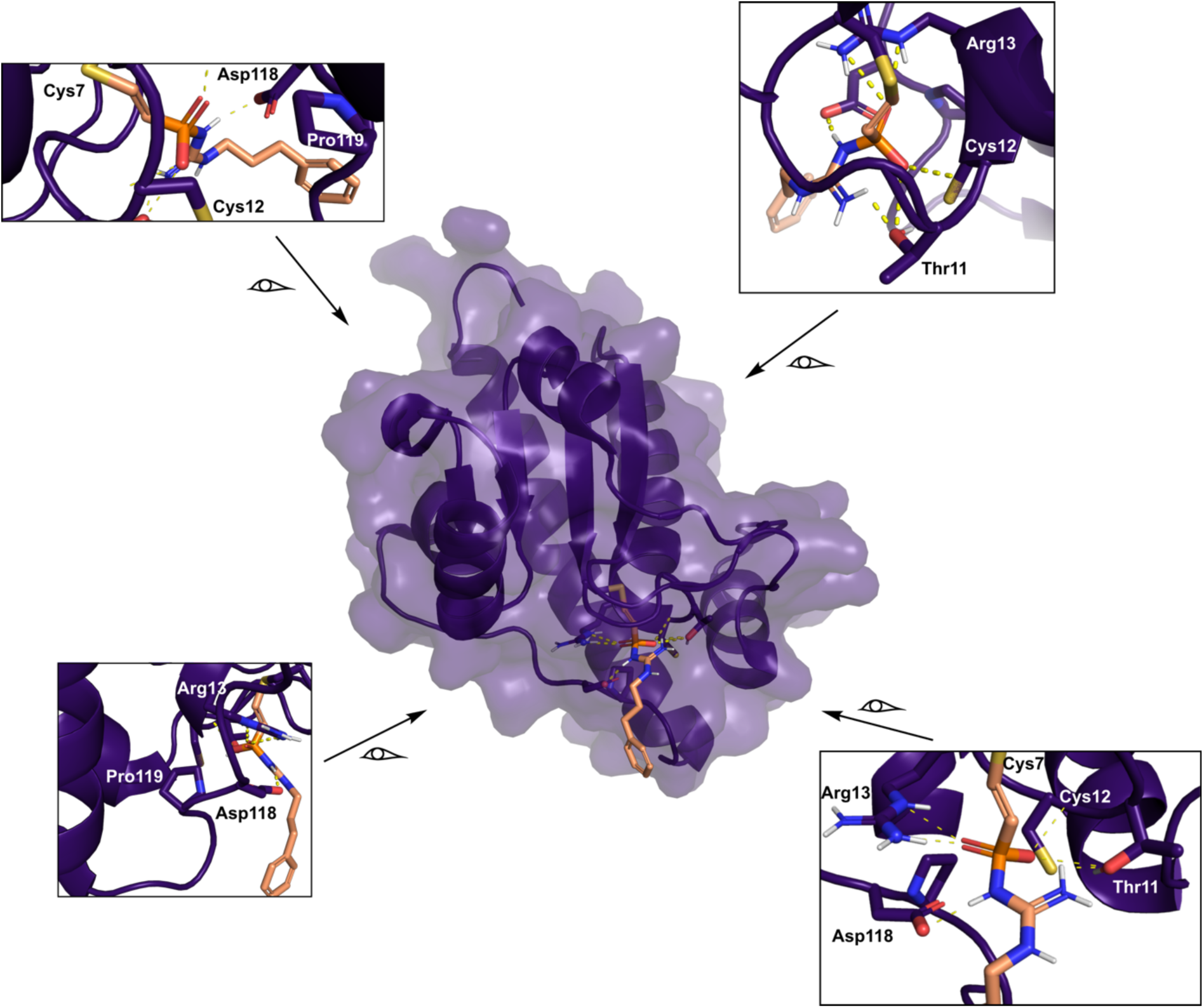
Covalent docking of P2b onto active site Cys7 of YwlE. (PDB: 4KK3). Yellow dotted lines indicate putative hydrogen-bonds. No steric clashes were observed for this probe.

**Extended Data Fig 3:**
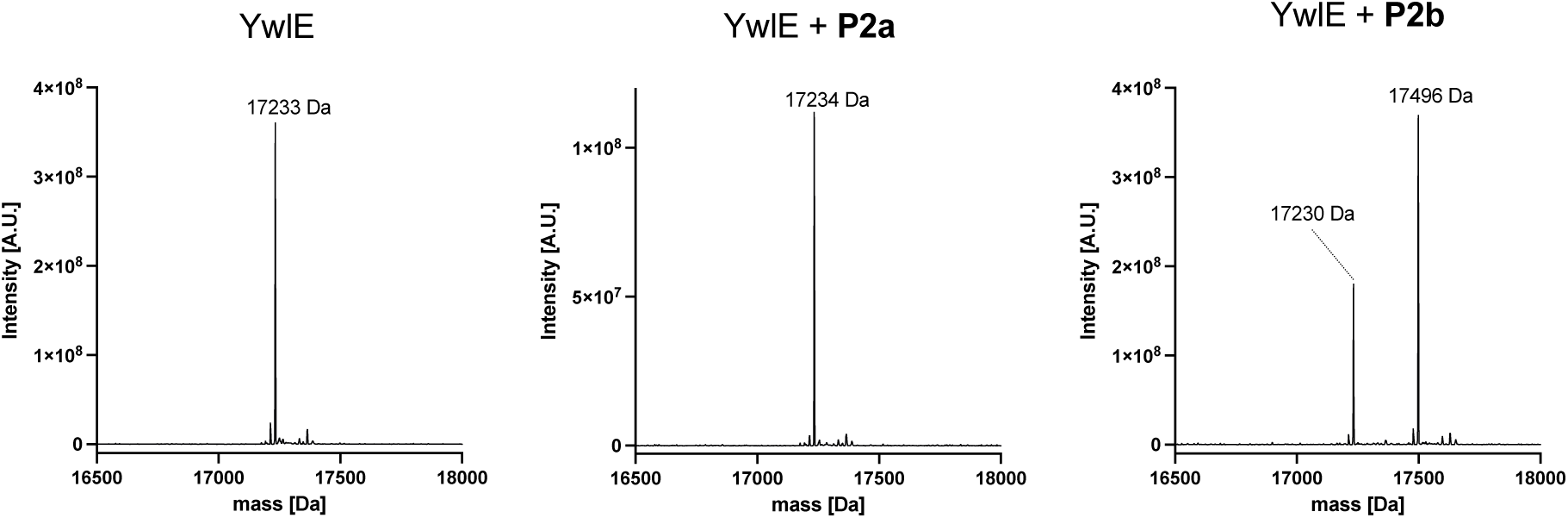
Intact-protein MS of recombinant YwlE_G.sp._ incubated with P2a/b. *Left panel: Deconvoluted intact MS spectra of YwlE_G.sp._* Middle panel: *Deconvoluted intact MS spectra of YwlE_G.sp._ after 24 h incubation with **P2a** and P2b (right panel). **P2a** does not show any modification, while **P2b** indicates significant modification of YwlE.* YwlE + **P2b** calc.: 17498 Da, found.: 17499 Da.

**Extended Data Fig 4:**
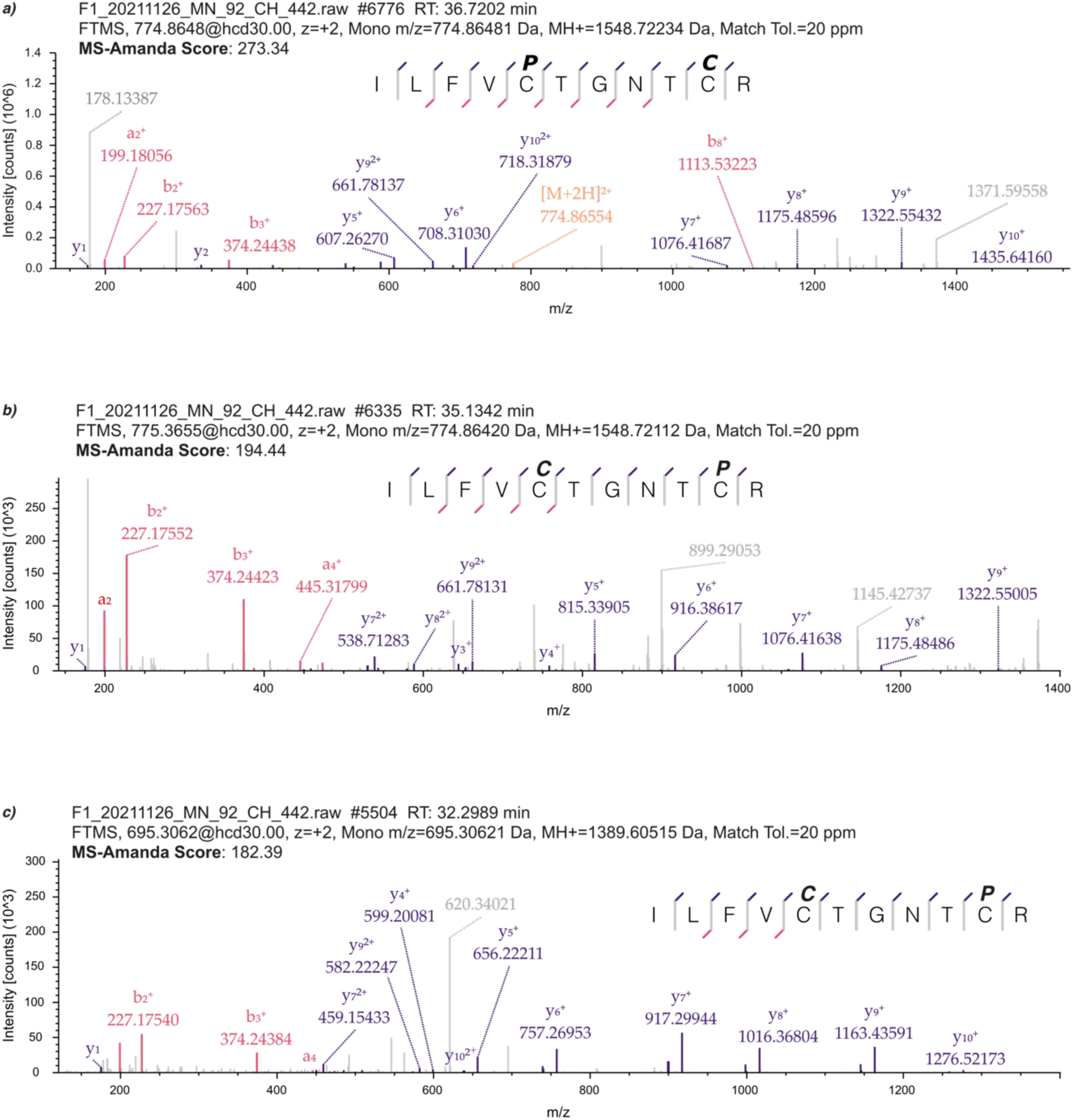
MS/MS analysis of *YwlE_G.sp._* labeled with P2b. a) overall best scoring spectrum indicating labelling of active site Cys7. b)-c) Peptide spectrum matches indicating labelling of Cys12. C = Carbamidomethylation, P = modification with **P2b**.

**Extended Data Fig 5:**
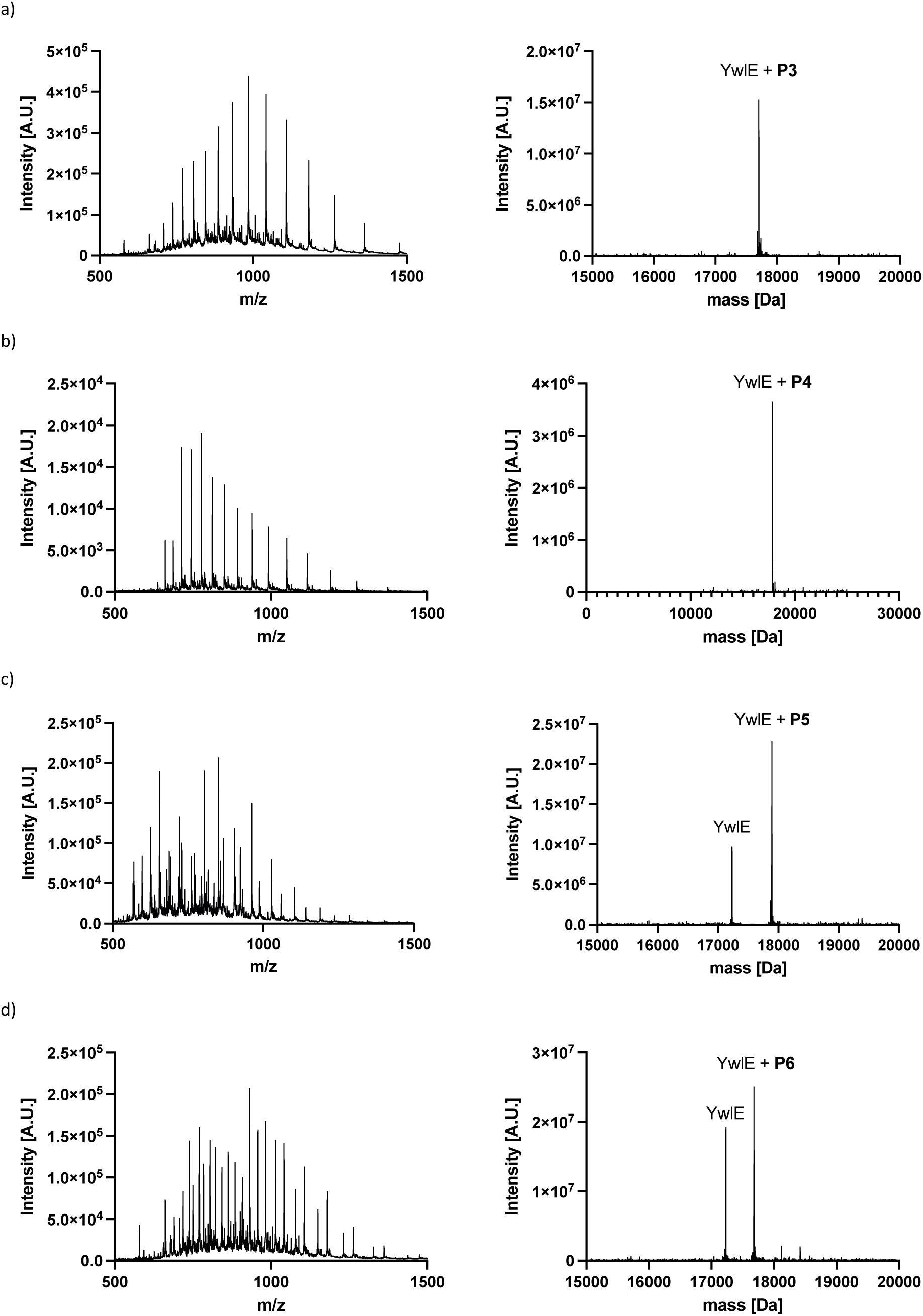
Intact protein MS analysis of *YwlE_G.sp._* labeled with probes P3-P6. a) Intact-protein MS analysis of YwlE_G.sp._ after reaction with biotin functionalized probe **P3** for 24 h. YwlE + **P3** calc.: 17706 Da, found.: 17705 Da. b) Intact-protein MS analysis of YwlE_G.sp._ after reaction with fluorescein functionalized probe **P4** for 24 h. YwlE + **P4** calc.: 17838 Da, found.: 17838 Da. c) Intact-protein MS analysis of YwlE_G.sp._ after reaction with TAMRA functionalized probe **P5** for 24 h. YwlE + **P5** calc.: 17892 Da, found.: 17893 Da. d) Intact-protein MS analysis of YwlE_G.sp._ after reaction with tetrazine functionalized probe **P6** for 24 h. YwlE + **P6** calc.: 17679 Da, found.: 17680 Da.

**Extended Data Fig 6:**
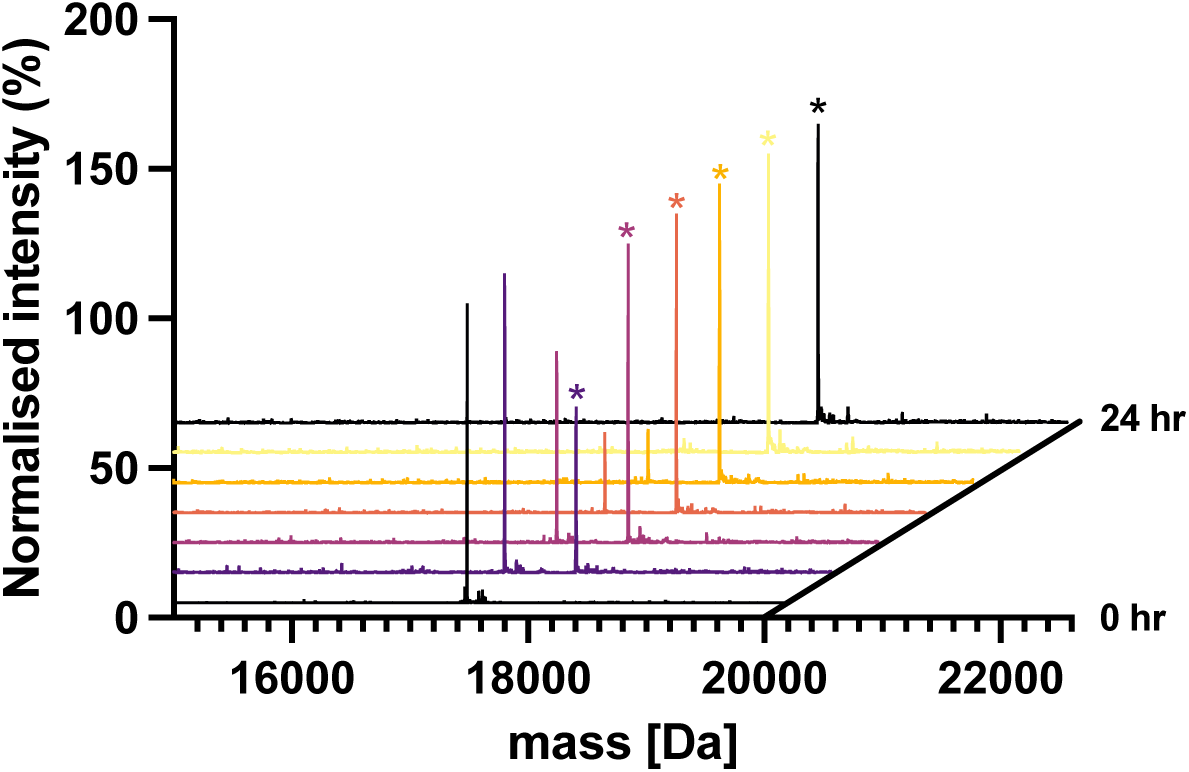
**Labelling Time-Course of *YwlE_G.sp._ using probe P4.*** Time-dependent labelling of *YwlE_G.sp._* (10 µM) using the fluorescein-functionalized pArg phosphatase probe **P4** (25 µM). The peak corresponding to probe functionalized YwlE is marked with an asterisk. Aliquots were measured at 0, 1, 2, 4, 8, 16 and 24 h, respectively.

**Extended Data Fig 7:**
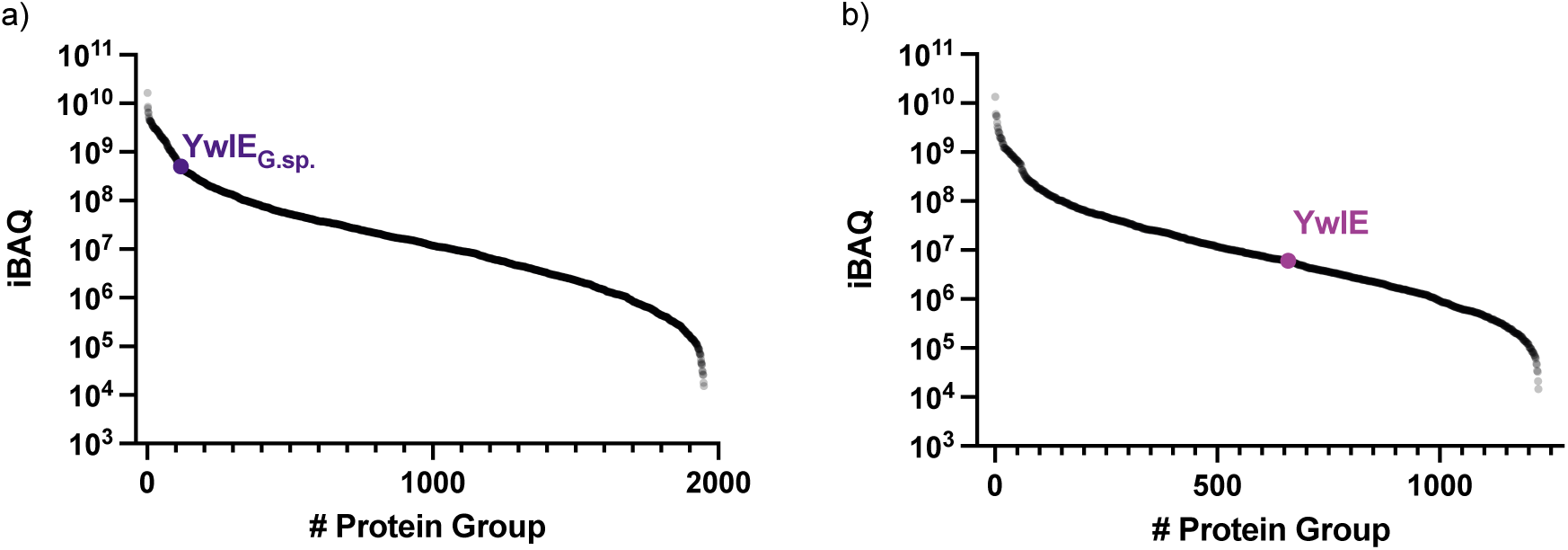
iBAQ based quantification of trypsin digested *E. coli* BL21 (a) or *B. subtilis* lysate (b).

**Extended Data Fig 8:**
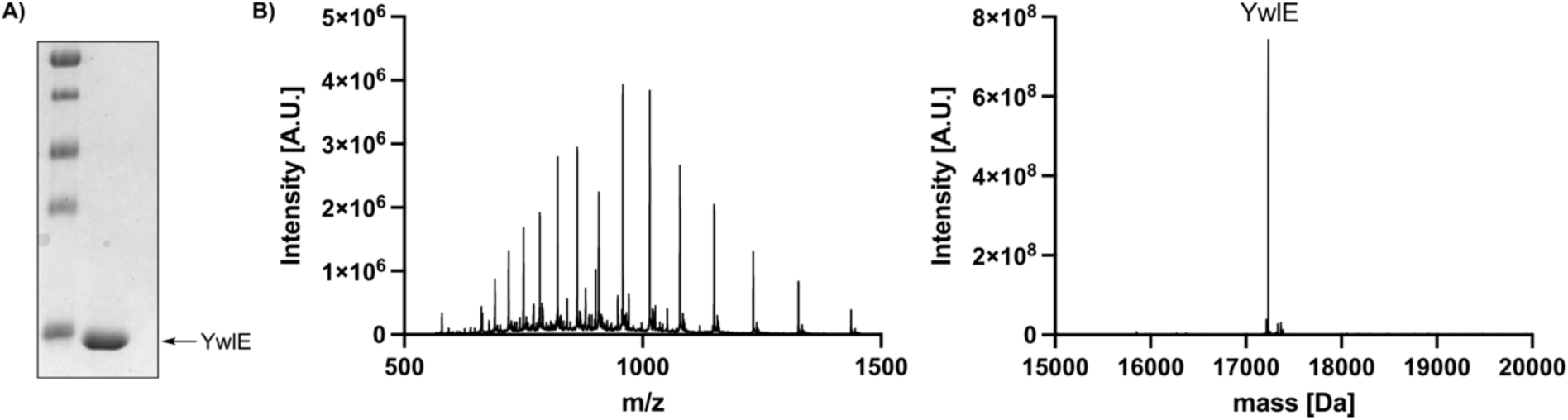
**Characterization of purified YwlE_G.sp._** A) SDS-PAGE analysis of the purified protein. B) intact-protein MS analysis of spectrum of YwlE_G.sp._, calc.: 17233 Da, found.: 17234 Da.

**Extended Data Fig 9:**
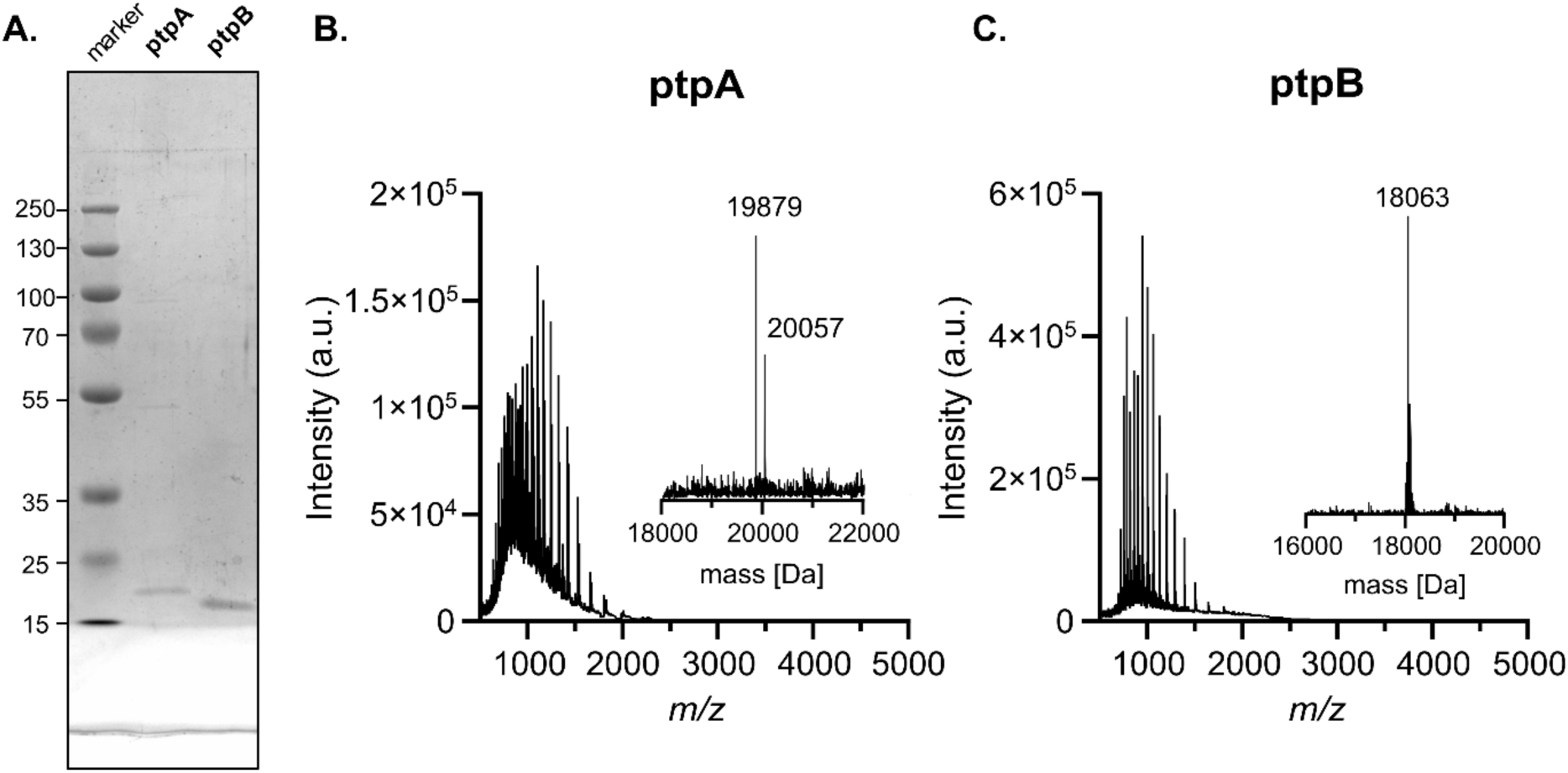
Characterization of purified PtpA and PtpB. A) SDS-PAGE analysis of the purified proteins. HRMS spectrum of B) PtpA, calc.: 19881 Da, [M+gluconylation] calc.: 20059, Da exp.: 19879 Da, 20057 Da, and C) PtpB calc.: 18063 Da, exp.: 18063 Da.

## Notes

### Competing Interest Statement

The authors have declared no competing interest.

