## Supplementary Information for "Active Site-Directed Probes for targeting Bacterial Phosphoarginine Phosphatases"

[1] Leibniz-Forschungsinstitut für Molekulare Pharmakologie (FMP), Robert-Rössle-Strasse 10, 13125 Berlin, Germany.

[2] Department of Chemistry, Humboldt Universität zu Berlin, Brook-Taylor-Strasse 2, 12489 Berlin, Germany.

[3]Department of Bioscience, Center for Functional Protein Assemblies (CPA), Technical University of Munich, Ernst-Otto-Fischer Strasse 8, 85748 Garching bei München, Germany.
*corresponding authors

### 1. Mass Spectrometry based ABPP

#### 1.1 Lysate Labelling and Sample Preparation

***E. coli* transfected with YwlE$_{G.sp.}$:**

Lysates (approx. 100 µg of protein each) were incubated with the indicated concentration of biotin-pArg-phosphatase-probe **P3** for 1 h at room temperature. Afterwards, proteins were precipitated with the 4-fold volume of acetonitrile and cetrifuged (14'00 rcf, 5 min). The solvent was removed and precipitates were resuspended in 80% EtOH by water bath sonication, followed by another round of centrifugation (10'000 rcf, 5 min). This procedure was repeated three times. The pellets were aspirated and left to dry on air for 5-10 minutes. Afterwards, the precipitate was redissolved in 800 µl 8 M urea (100 mM Tris, pH 8) followed by the addition of 20 µl prewashed streptavidin coated agarose beads (Pierce™ Streptavidin Agarose, 20347). After 2 h of incubation at room temperature on a rotor, the supernatant was removed and beads were washed as follows: 3x 8M Urea, 1x 1M NaCl in PBS, 3x 1% SDS in PBS (each 1 ml). Collected beads were resuspended in 20 µl 1x Lammli-buffer containing 1% SDS and heated to 95 °C for 10 miutes. The hot supernatant of the beads was loaded in a 12% SDS gel and seperated partially (approx. 1 cm). Subsequently the samples were processed via in gel trypsin digestion. In brief, the gel was coomassie stained and corresponding lanes were cut out and collected into 1.5 ml eppendorf tubes. Gel pieces were destained by alternate washing with 50 mM ABC buffer (pH 8.5) and ABC buffer/MeCN (1:1), respectively. Destained gels were shrinked by the addition of 100% MeCN and left briefly on air to dry. The pieces were rehydrated in 50 mM ABC buffer containing 10 mM DTT and incubated at 56 °C for 30 minutes. Subsequently, 20 mM iodoacetamide were added and the solution was incubated for 45 minutes at room temperature in the dark. After alkylation was comlete, the solvent was removed and the gel pieces were again dehydrated by the addtion of 100% MeCN. Dried gel pieces were rehydrated in 50 mM ABC buffer containing 10 ng trypsin/µl (Promega, V5111) at 4 °C for approx. 15 minutes. The excess solvent was removed and 20 µl of additional 50 mM ABC buffer containing 10 ng trypsin/µl were added followed by over night incubation at 37 °C. The next day, the mixture was acidified and the supernatant was transferred to a different tube. Gel pieces were once more extracted with water/MeCN (1:1) containing 1% formic acid. Combined supernatants were evaporated to dryness and resuspended in MQ water containing 1% MeCN and 0.1% TFA.

***B. subtilis*** (DSM347):

Lysates (approx. 200 µg of protein each) were incubated with the indicated concentration of biotin-pArg-phosphatase-probe **P3** for 1 h at room temperature. Afterwards, the samples were processed according to a procedure published by Becker et al.[1]: A total of 50 µl of mixed hydrophobic and hydrophilic carboxylate-coated magnetic beads (1:1, prewashed with PBS; Seramag™, Cytavia) was added to the click reaction mixture followed by 600 µl of absolute ethanol to precipitate the proteins. Beads were washed thrice with of 80% ethanol and proteins were eluted by two sequential washes of 0.2% SDS in PBS. The eluted proteins were directly transferred onto 30 µl of streptavidin-coated magnetic beads (prewashed with 0.2% SDS in PBS; New England Biolabs) and incubated on a shaker for 1 h at room temperature. Streptavidin-beads were washed thrice with 8 M urea, PBS pH 7.4 and MilliQ water. Washed beads were resuspended in 100 mM ABC buffer, followed by reduction and alkylation (5 mM TCEP, 10 mM iodoacetamide, 37 °C, 30 min) and proteins were trypsinized overnight at 37 °C (1.5 µg/sample, Promega). The supernatant was removed, acidified to 1% TFA, centrifuged at 10'000 rcf for 5 minutes and stored at -20 °C until measurement.

***S. aureus*** (USA3000), ***L. monocytogenes*** (EGD-e), ***M. smegmatis*** (DSM43756):
Preparative MS-based ABPP was performed as described before.[2] All reagents used were of LC-MS grade. Sample processing followed an adapted protocol for magnetic bead enrichment. To each sample, 10 µL of a 2-fold concentrated 1:1 mix of washed (3x with $H_2O$) hydrophobic and hydrophilic carboxylate-coated magnetic beads (Cytiva, cat#65152105050250 and 45152105050250) was added. This was followed by the addition of 175 µL EtOH to precipitate the proteins onto the beads. The subsequent steps were carried out using an automated liquid handling system (Hamilton Microlab Prep, Hamilton). The plate was then incubated for 5 min at 500 rpm and r.t., following washing of the beads. Each washing step was performed with the help of a 96-well ring magnet (Alpaqua, Magnum FLX). For this, the plate was placed on it, and the supernatant was removed slowly (20 µL/s) to avoid removing any beads. Washing involved removing the plate from the magnet, adding the respective washing solution, and shaking for 1 min at 800 rpm and r.t. The samples were washed three times with 180 µL of 80% EtOH and once with 180 µL acetonitrile. Proteins were eluted from the beads by adding 75 µL of 0.2% SDS in PBS, followed by 5 min of incubation at 800 rpm and 40 °C. The plate was placed back on the magnet to transfer the supernatant into new wells. This elution step was repeated, resulting in a total volume of 150 µL of eluted proteins. Next, to each well containing the eluted protein samples, 50 µL of washed (3x with 0.2% SDS in PBS) streptavidin magnetic beads (New England Biolabs, cat#25S1420S) were added. The plate was sealed and incubated for 1 h at 800 rpm and r.t. in a plate shaker with a heated lid (ThermoMixer C, Eppendorf) to allow binding of the labeled proteins to the streptavidin beads. After this incubation step, the plate was further processed with the liquid handling system. Washing of the beads was done three times with 180 µL of 0.1% NP-40 in PBS, two times with 180 µL of 6 M urea, and three times with 200 µL $H_2O$. Protein digestion was performed with 1 µL trypsin (trypsin/protein ratio 1:100, 0.5 µg/µL, sequencing grade, Promega) in 100 µL of 50 mM TEAB overnight at 800 rpm and 37 °C in a plate shaker with a heated lid. The plate was tightly sealed during digestion. The next day, peptides were eluted from the beads in the liquid handling system with 50 µL of 3% FA and transferred into new wells for desalting. Desalting was performed using pre-equilibrated stage tips containing two layers of styrenedivinylbenzene-reverse phase sulfonate (SDB-RPS) disks (Empore, 3M). The stage tips were equilibrated with 150 µL wash buffer 1 (1% TFA in isopropanol) before loading the samples. Samples were loaded for 10 min at 500 x g, followed by washing with 150 µL wash buffer 1 for 10 min at 800 x g, and another wash with 150 µL wash buffer 2 (0.2% TFA in $H_2O$). Peptides were eluted with 50 µL elution buffer (1% ammonia, 80% acetonitrile) for 5 min at 300 x g, followed by 5 min at 800 x g. Eluted peptide samples were dried using a centrifugal evaporator (Concentrator Plus, Eppendorf), following reconstitution in 35 µL of 1% FA. Preparative MS samples were performed in n = 4 biologically independent replicates.

#### 1.2 Mass spectrometry analysis of pull-down samples

***E. coli*** transfected with $YwlE_{GS}$ & ***B. subtilis***:
Before MS/MS analysis a quality-control run was conducted on a Waters XEVO G2-XS QTof instrument to ensure complete digestion. Samples were diluted based on the QC-runs to ensure equal loading. LC-MS/MS analysis was performed using an UltiMate 3000 RSLC nano LC system coupled on-line to an Orbitrap Fusion mass spectrometer (Thermo Fisher Scientific). For sample loading a PepMap C-18 trap-column (Thermo Fischer Scientific) of 0.075 mm ID x 50 mm length, 3 µm particle size and 100 Å pore size was used. The loading mobile phase A contained 1% acetonitrile and 0.05% TFA acid in water, and mobile phase B 0.05% TFA acid in acetonitrile. Reversed-phase separation was performed using a 50 cm analytical column (in-house packed with Poroshell 120 EC-C18, 2.7µm, Agilent Technologies) with mobile phase A contained 0.1% formic acid in water, and mobile phase B 0.1% formic acid in

acetonitrile using a 93 minutes gradient (4-5%B 0-8 minutes; 5-25%B in 8-74 minutes; 25-28%B 74-80 minutes; 28-31%B 80-86 minutes; 31-36%B 86-92 minutes; 36-40%B 92-95 minutes; 40-50%B 95-96 minutes; 50-80%B 96-101 minutes; 80%B 101-104 minutes; 80-4%B 104-104.1 minutes). Data was acquired using survey scans in a range of 375 to 1500 m/z with a resolution of 120k and an AGC target value of 4e5. Precursor ions with charge states 2-4 were isolated with a mass selecting quadrupole (isolation window 1.6 m/z) with 40 sec dynamic exclusion (+/- 10 ppm). Precursor ions were fragmented using higher-energy collisional dissociation (HCD) applying a normalized collision energy (NCE) of 30. The maximum injection time was set to 22 ms to collect 5e4 precursor ions. Fragment ion spectra were acquired in the Orbitrap with a resolution of 15k (FWHM).

***S. aureus, L. monocytogenes***, ***M. smegmatis***:

Peptides were measured and online-separated using an *UltiMate 3000 nano* HPLC system (*Dionex, Sunnyvale, USA*) coupled to a *timsTOF Pro* (*Bruker, Billerica, USA*) mass spectrometer via a *CaptiveSpray* nano-electrospray ion source (*Bruker, Billerica, USA*) and *Sonation* (*Biberach, Germany*) column oven. Peptides were first loaded on the trap column (*Acclaim PepMap 100 C18*, 75 µm ID x 2 cm, 3 µm particle size, *Thermo Fisher Scientific, Waltham, USA*), washed with 0.1% formic acid in water for 7 min at 5 µL/min and subsequently transferred to the separation column (*Aurora* C18 column, 25 cm × 75 µm, 1.7 µm, *IonOpticks, Fitzroy, Australia*) and separated over a 60 min gradient from 5% to 28% B, then to 40% B over 13 min, followed by 10 min at 95% before re-equilibration and at a flow rate of 400 nL/min. The mobile phases A and B were 0.1% (v/v) formic acid in water and 0.1% (v/v) formic acid in acetonitrile, respectively.

The *timsTOF Pro* was operated in data-independent dia-PASEF mode with the dual TIMS analyzer operating at equal accumulation and ramp times of 100 ms each with a set 1/K0 ion mobility range from 0.60 to 1.60 V × s × $cm^{-2}$. for MS1 scans. The dia-PASEF settings for fragmentation were set to a mass range of 400 to 1201 m/z and an ion mobility range of 0.60 to 1.43 V × s × $cm^{-2}$. Two ion mobility isolation windows were performed per dia-PASEF scan with 26 m/z window widths. A total of 32 isolation windows with 1 m/z overlaps to cover the mass range were used resulting in 16 dia-PASEF scans per MS1 scan and an estimated total cycle time of 1.80 s (see table 1). The collision energy was ramped linearly as a function of the mobility from 59 eV at 1/K0 = 1.3 V × s × $cm^{-2}$ to 20 eV at 1/K0 = 0.85 V × s × $cm^{-2}$. TIMS elution voltages were calibrated linearly to obtain the reduced ion mobility coefficients (1/K0) using three Agilent ESI-L Tuning Mix ions (m/z 622, 922 and 1,222) spiked on the CaptiveSpray Source inlet filter.

***Table S1:*** *DiaPASEF – long gradient method: isolation window layout in detail.*

| MS Type | Scan | Start IM [1/K0] | End IM [1/K0] | Start Mass [m/z] | End Mass [m/z] |
|---|---|---|---|---|---|
| MS1 | 0 | 0.6 | 1.6 | 100 | 1700 |
| dia-PASEF | 1 | 0.9 | 1.2 | 800 | 826 |
| dia-PASEF | 1 | 0.6 | 0.9 | 400 | 426 |
| dia-PASEF | 2 | 0.92 | 1.22 | 825 | 851 |
| dia-PASEF | 2 | 0.62 | 0.92 | 425 | 451 |
| dia-PASEF | 3 | 0.93 | 1.23 | 850 | 876 |
| dia-PASEF | 3 | 0.63 | 0.93 | 450 | 476 |
| dia-PASEF | 4 | 0.95 | 1.25 | 875 | 901 |

| | | | | | |
|---|---|---|---|---|---|
| dia-PASEF | 4 | 0.65 | 0.95 | 475 | 501 |
| dia-PASEF | 5 | 0.96 | 1.26 | 900 | 926 |
| dia-PASEF | 5 | 0.66 | 0.96 | 500 | 526 |
| dia-PASEF | 6 | 0.98 | 1.28 | 925 | 951 |
| dia-PASEF | 6 | 0.68 | 0.98 | 525 | 551 |
| dia-PASEF | 7 | 0.99 | 1.29 | 950 | 976 |
| dia-PASEF | 7 | 0.69 | 0.99 | 550 | 576 |
| dia-PASEF | 8 | 1.01 | 1.31 | 975 | 1001 |
| dia-PASEF | 8 | 0.71 | 1.01 | 575 | 601 |
| dia-PASEF | 9 | 1.02 | 1.32 | 1000 | 1026 |
| dia-PASEF | 9 | 0.72 | 1.02 | 600 | 626 |
| dia-PASEF | 10 | 1.04 | 1.34 | 1025 | 1051 |
| dia-PASEF | 10 | 0.74 | 1.04 | 625 | 651 |
| dia-PASEF | 11 | 1.06 | 1.36 | 1050 | 1076 |
| dia-PASEF | 11 | 0.76 | 1.06 | 650 | 676 |
| dia-PASEF | 12 | 1.07 | 1.37 | 1075 | 1101 |
| dia-PASEF | 12 | 0.77 | 1.07 | 675 | 701 |
| dia-PASEF | 13 | 1.09 | 1.39 | 1100 | 1126 |
| dia-PASEF | 13 | 0.79 | 1.09 | 700 | 726 |
| dia-PASEF | 14 | 1.1 | 1.4 | 1125 | 1151 |
| dia-PASEF | 14 | 0.8 | 1.1 | 725 | 751 |
| dia-PASEF | 15 | 1.12 | 1.42 | 1150 | 1176 |
| dia-PASEF | 15 | 0.82 | 1.12 | 750 | 776 |
| dia-PASEF | 16 | 1.13 | 1.43 | 1175 | 1201 |
| dia-PASEF | 16 | 0.83 | 1.13 | 775 | 801 |

#### 1.3 Data Analysis of Pull-Down Experiments

***E. coli* transfected with YwlE$_{GS}$ & *B. subtilis***:
The obtained raw-data was analyzed using FragPipe (v20 to 21) using the built-in LFQ-MBR workflow using the whole human proteome as search space and applying the following MSFragger[3] settings: Precursor mass tolerance: +/-10 ppm; Fragment mass tolerance: +/-20 ppm; Mass calibration & parameter optimization enabled; Isotope error: 0/1/2; Enzyme: Trypsin (cuts after K & R, no cut before P), Peptide length: 7-50 AA; Peptide mass range: 500-5000 Da; Variable modifications: Oxidation (M, 15.9949 Da, up to 3x), Acetylation (N-term, 42.0106 Da); Carbamidomethylation of C was set as fixed modification (57.02146 Da). Validation was performed via Percolator & ProteinProphet and an FDR of

1% was applied. IonQuant[4] was used for label free quantification, with MBR (FDR of 1%) and normalization across runs enabled. Analyzed files were further processed using FragPipe-Analyst[5] as follows: Proteins have to be quantified in >65% of the files for at least one condition and in >66% of the files globally. Variance stabilization was enabled, and Benjamin Hochberg type FDR correction was enabled.
Unless stated differently the following significance cut offs were used: $\log_2$ fold-change >2; $-\log_{10}$ (adjusted) p-value >2.

***S. aureus*, *L. monocytogenes*, *M. smegmatis***:
DIA-NN version 1.8.1 was used with "FASTA digest for library-free search" and "deep learning-based spectra, RTs and IMs prediction". "Trypsin/P" with a maximum of two missed cleavages was set as protease. The maximum number of variable modifications was set to 0 and N-terminal methionine excision and cysteine carbamidomethylation selected as fixed modifications. Peptide precursors with a length of 7 to 30 amino acids and a charge range from 2 to 4 were considered. The precursor m/z range was set from 300 to 1800 m/z, the fragment ion ranges from 200 to 1800 m/z. "Generate spectral library" and quantities matrices were activated and a precusor FDR of 1% was used. Mass accuracy, MS1 accuracy and scan window was set to automatic (0), "use isotopologues", match between runs (MBR) and "remove likely interferences" were activated. The neural network classifier was used in "single-pass mode", proteins were inferred from genes and the quantification strategy set to "Robust LC (high precision)". For cross-run normalisation "RT-dependent" was selected and "Smart profiling" used for library generation. The additional option "--relaxed-prot-inf" was used. The protein group quantity matrix was used for further downstream analysis in Perseus. The respective FASTA-files were obtained from UniProt (*S. aureus* USA300, Proteome ID: UP000000793; *L. monocytogenes* EGD-e Proteome ID: UP000000817; *M. smegmatis* DSM43756, Proteome ID: UP000000757). Perseus 2.0.9.0[6] was used for the downstream analysis. Protein-groups text files from the DIA-NN 1.8 analysis were loaded into Perseus and LFQ intensities were $\log_2$ transformed. Subsequently rows were annotated in groups and potential contaminants were removed. Missing value imputation was performed over the total matrix and to visualize the data a scatterplot was created based on two-sided two sample Student's t-test (FDR = 0.05).

### 2. Synthetic Procedures

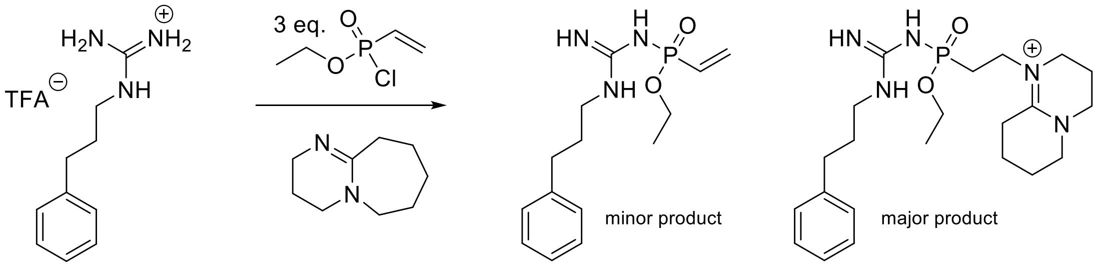

*Scheme S1: Initial attempt towards probe **P1a**. DBU is able to deprotonate guanidinium **1** to facilitate reaction with ethyl vinylphosphonochloridate (**2**). However, DBU is sufficiently nucleophilic to undergo a subsequent addition reaction to the vinyl group, resulting in unwanted side reactions.*

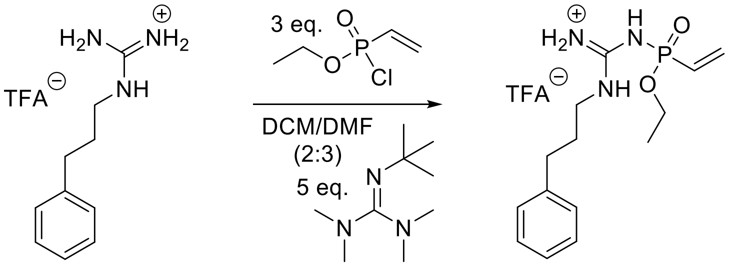

*Scheme S2: Optimized synthetic route towards the synthesis of vinyl-guanidin-phosphonamidates from chloro-vinyl-phosphonates.*

#### 1-(3-phenylpropyl) guanidine (**1**):

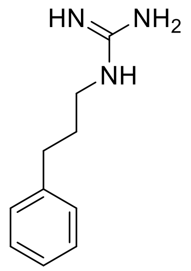

3-phenyl-1propylamine (136 mg, 1 mmol, 1 eq.) was dissolved in 5 ml of a 1 M aq. NaOH-solution and 5 ml acetonitrile. 1.2 eq. S-methylisothiourea hemisulfate (167 g, 1.2 mmol) were added in portions and the reaction was stirred for 24 h at room temperature. The solvent was evaporated under reduced pressure and the residue purified via HPLC to yield compound **1** as a white powder. (TFA salt, 55% yield)

**$^1$H NMR** (300 MHz, $D_2O$): δ 7.34 – 7.11 (m, 3H), 3.06 (t, J = 6.9 Hz, 1H), 2.60 (t, J = 7.5 Hz, 1H), 1.89 – 1.73 (m, 1H)
**$^{13}$C-NMR** (75 MHz, $D_2O$) δ 156.61, 141.50, 128.65 (2C), 128.47 (2C), 126.16, 40.40, 31.91, 29.38.
**HRMS** for $C_{10}H_{16}N_3^+$ $[M+H]^+$ calc.: 178.1339 Da; found: 178.1345 Da.

ethyl vinylphosphonochloridate (**2**):

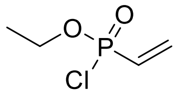

1.64 g of diethyl vinylphosphonate were dissolved in 10 ml dry DCM. Over the course of 5 minutes, 2 eq. oxalylchloride were added dropwise, followed by 16 h reaction at room temperature. Afterwards, the mixture was refluxed for 1 h and the conversion of the starting material was monitored by $^{31}$P-NMR. After full consumption of the starting material, all volatiles were removed and the residue was dissolved in 10 ml dry DCM to form an approx. 1 M solution of **2** that was used without further purification.

**$^{31}$P NMR** (122 MHz, ) δ 25.76.

ethyl N-(N-(3-phenylpropyl)carbamimidoyl)-P-vinylphosphonamidate (**P1a**):

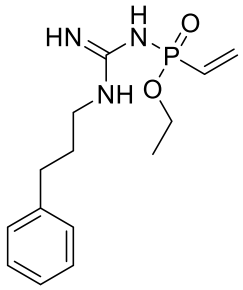

36 mg **1** (0.2 mmol, 1 eq.) was dissolved in 2 ml dry DMF under argon and 5 eq. Barton's base were added. Subsequently, 3 eq. of freshly prepared **2** (1 M in DCM) were added dropwise while the solution was stirred on ice. After another 30 minutes of stirring at room temperature, the solvent was removed under reduced pressure and the residue purified via preparative HPLC. Compound **3** was obtained as a white solid. (TFA salt, 42 mg, 71% yield)

**$^{1}$H NMR** (600 MHz, Methanol-$d_4$) δ 7.30 (t, $J$ = 7.5 Hz, 2H), 7.26 – 7.16 (m, 3H), 6.51 – 6.26 (m, 3H), 4.35 – 4.17 (m, 2H), 3.29 (t, $J$ = 7.0 Hz, 2H), 2.76 – 2.66 (m, 2H), 1.97 (p, $J$ = 7.2 Hz, 2H), 1.41 (t, $J$ = 7.1 Hz, 3H).
**$^{13}$C NMR** (151 MHz, Methanol-$d_4$) δ 140.61, 136.78, 128.19, 128.02, 125.85, 125.63 (d, $J$ = 175.0 Hz), 62.88 (d, $J$ = 6.6 Hz), 40.99, 32.17, 29.55, 15.07 (d, $J$ = 6.4 Hz).
**$^{31}$P NMR** (243 MHz, Methanol-$d_4$) δ 16.60.
**HRMS** for $C_{14}H_{23}N_3O_2P^+$ $[M+H]^+$ calc.: 296.1522 Da; found: 296.1533 Da

#### Synthetic attempts towards P1b:

Initial attempts towards **P1b** envisioned the direct activation of diethyl-ethynyl phosphonate using oxalylchloride analogously to the corresponding vinyl-compound (Scheme S3a). However, even large excess (>20 eq.), catalytic amounts of DMF and prolonged reflux in pure oxalylchloride did not result in the desired conversion to the phosphonochloridate. Therefore, we thought to first hydrolyze diethyl-ethynyl phosphonate and perform the activation in a separate step (Scheme S3b). However, diethyl-ethynyl phosphonate also proved to be extremely stable towards basic hydrolysis. In a further attempt we started from ethynyl-phosphonamidates that can be hydrolyzed under acidic conditions (Scheme S3c). To our delight, the acid mediated hydrolysis proceeded to completion within less than 1 h, but we were unsuccessful in isolating the phosphonic acid derivative from the aqueous phase. (Interestingly, we observed a similar phenomenon later using ortho-nitrobenzyl functionalized phosphonates).

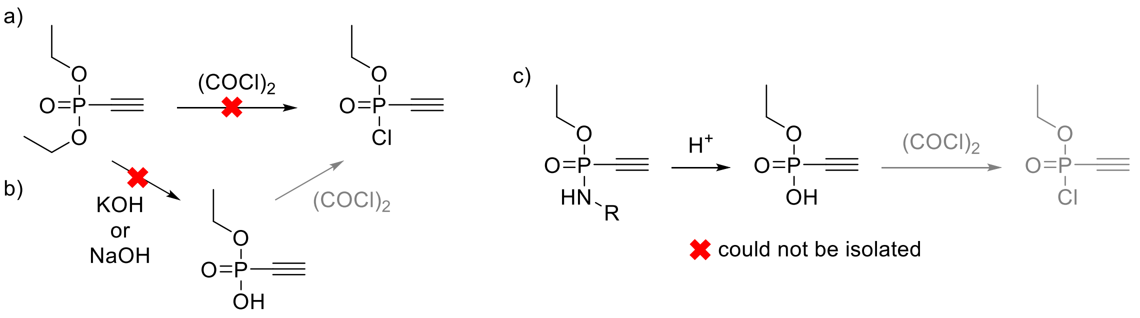

*Scheme S3: Unsuccessful attempts towards* ***P1b***

Finally, equipping the ethynyl-group with a triisopropyl-protecting group added sufficient hydrophilicity to transfer the phosphonic acid to the organic phase (Scheme S4).

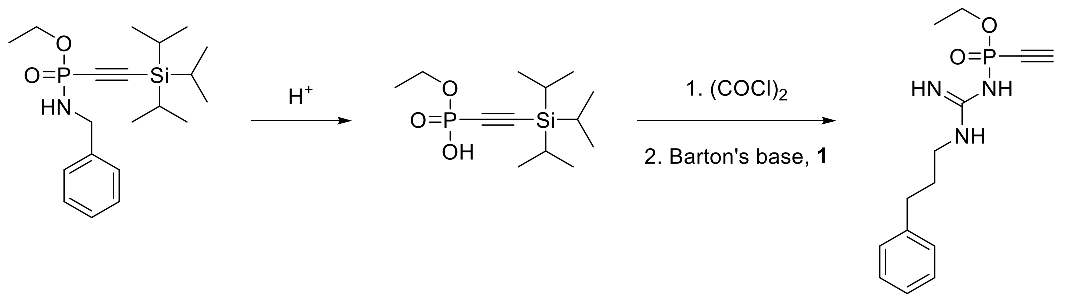

*Scheme S4: Synthetic route that facilitated the synthesis of probe* ***P1b***

ethyl *N*-benzyl-*P*-((triisopropylsilyl)ethynyl)phosphonamidite (**3**):

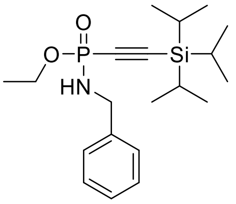

TIPS acetylene (5 mmol) was dissolved in 3 ml absolute THF under argon and cooled to -78 °C. 1 eq. of n-BuLi (2.5 M in Hexanes) was added dropwise, the misture was allowed to warm to r.t. and stirred for 30 minutes. After cooling the reaction to -78 °C, 1 eq. of diethyl-chloro-phosphite was added dropwise and the mixture was allowed to warm to r.t.. A 1 mmol aliquote of the intermediate was reacted with benzyl-azide overnight at 40 °C followed by the addition of $H_2O$. The reaction was vigorously stirred for two hours and organics were extracted using EtOAc. Combined organic layers were dried over $MgSO_4$, filtered, evaporated and purified via column chromatography on silica gel (Hexane/EtOAc; 1:1) to obtain **3**.

**$^1$H NMR** (300 MHz, Chloroform-*d*) δ 7.46 – 7.23 (m, 5H), 4.28 – 4.06 (m, 4H), 1.38 (td, *J* = 7.1, 1.6 Hz, 3H), 1.11 (d, *J* = 3.0 Hz, 21H).
**$^{31}$P NMR** (122 MHz, Chloroform-*d*) δ -2.83.

#### ethyl hydrogen ((triisopropylsilyl)ethynyl)phosphonate (**4**):

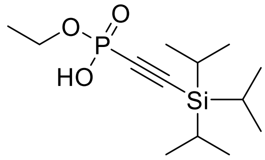

Compound **3** was dissolved in 95/5 TFA/$H_2O$ and upon complete cleavage, the solvent was blown off and phosphonic acid **4** was isolated by extraction from 1 M HCl/EtOAc. (95% yield)

**$^1$H NMR** (600 MHz, Chloroform-*d*) δ 11.19 (s, 1H), 4.17 – 4.02 (m, 2H), 1.34 (t, *J* = 7.1 Hz, 3H), 1.10 (h, *J* = 4.2, 3.6 Hz, 21H).
**$^{13}$C NMR** (151 MHz, Chloroform-*d*) δ 105.29 (d, *J* = 40.2 Hz), 97.51 (d, *J* = 278.2 Hz), 62.96 (d, *J* = 5.2 Hz), 18.33, 15.89 (d, *J* = 7.3 Hz), 10.85.
**$^{31}$P NMR** (243 MHz, Chloroform-*d*) δ -8.45.
**HRMS** for $C_{12}H_{28}O_3PSi^+$ $[M+H]^+$ calc.: 291.1540 Da; found: 291.1519 Da

#### ethyl P-ethynyl-N-(N-(3-phenylpropyl)carbamimidoyl)phosphonamidate (**P1b**):

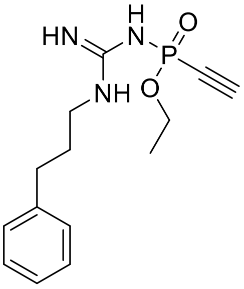

Compound **4** was dissolved in 1 ml oxalyl chloride under argon and 1 µl dry DMF was added. The reaction mixture was heated to reflux for 1 h. Volatiles were evaporated and co-evaporated two additional times with 1 ml DCM. Meanwhile compound **1** was dissolved in dry DMF, 5 eq. Barton's base were added, and the mixture was added to the activated $P^V$ compound. After consumption of the starting material the product was purified by semi-preparative HPLC. (6 mg, 41% yield)

**$^1$H NMR** (300 MHz, DMSO-*d*$_6$+TFA**)** δ 7.30 – 7.12 (m, 7H), 7.09 (s, 1H), 6.92 (s, 1H), 4.77 (d, *J* = 13.8 Hz, 1H), 4.21 (dqd, *J* = 9.6, 7.1, 2.9 Hz, 2H), 3.23 (q, *J* = 6.7 Hz, 2H), 2.67 – 2.57 (m, 2H), 1.90 – 1.72 (m, 2H), 1.31 (t, *J* = 7.0 Hz, 3H).
**$^{31}$P NMR** (122 MHz, DMSO-*d*$_6$+TFA**)** δ -11.95.

**Synthetic attempts towards P2a:**

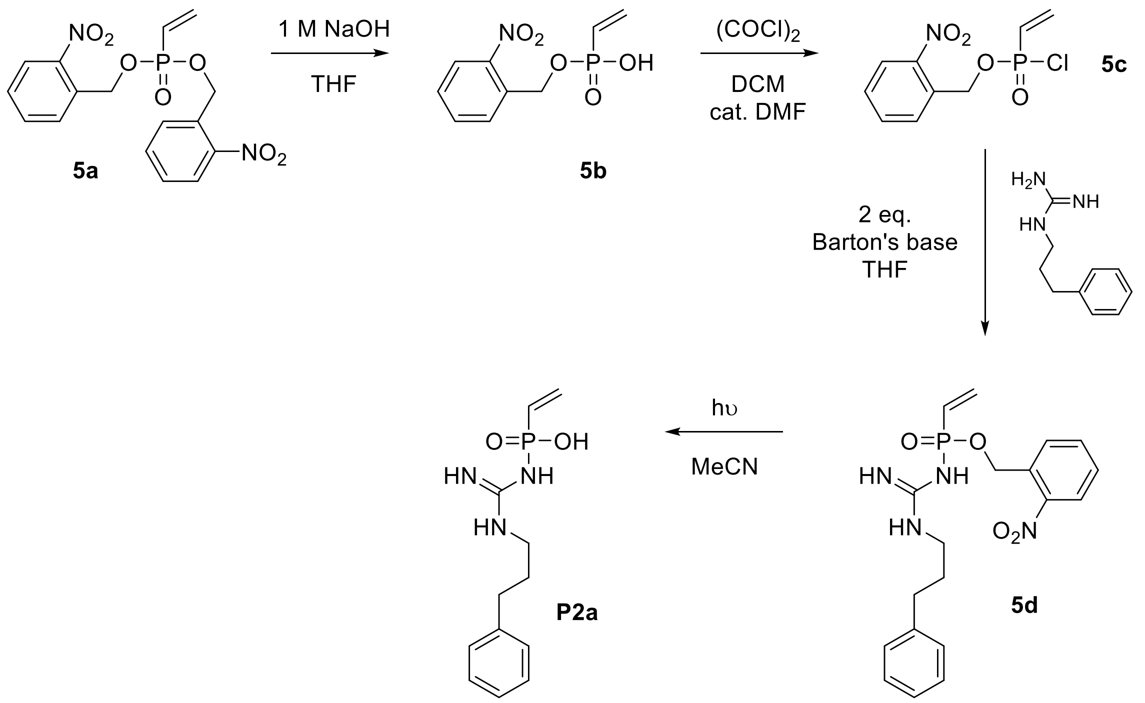

*Scheme S5: Synthetic route towards the putative vinyl-$P^V$ pArg-phosphatase probe* ***P2a***

bis(2-nitrobenzyl) vinylphosphonate (**5a**):

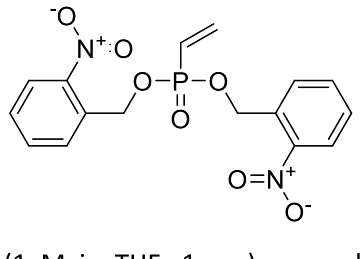

A 250-ml Schlenk flask was charged with 440 µl $PCl_3$ (5 mmol, 1 eq.) in 20 ml of dry $Et_2O$ under an argon atmosphere, cooled to -78 °C and 2 eq. *o*-nitro-benzyl alcohol (0.1 M in $Et_2O$ containing 2 eq. pyridine) were added dropwise. The solution was allowed to warm to room temperature and stirred for 30 minutes during which a white precipitate formed. To this solution, 5 ml of vinylmagnesium bromide (1 M in THF, 1 eq.) were slowly added and the reaction was again allowed to warm to room temperature. After another 30 minutes of stirring at room temperature, 2 eq. Luperox were added (1.5 ml 70% in water). After the oxidation was complete, the reaction was extracted 3x with 50 ml 1 M aq. HCl. Combined organic layers were dried over $MgSO_4$, filtered, evaporated and purified via column chromatography on silica gel (Hexane/EtOAc; 1:1) to obtain 699 mg of 5**a** as an off-white solid (37% yield).

$^{1}$**H NMR** (600 MHz, Chloroform-*d*) δ 8.13 (dd, *J* = 8.2, 1.2 Hz, 2H), 7.82 – 7.76 (m, 2H), 7.73 – 7.67 (m, 2H), 7.55 – 7.49 (m, 2H), 6.58 – 6.10 (m, 3H), 5.53 (dd, *J* = 7.6, 2.8 Hz, 4H).
$^{13}$**C NMR** (151 MHz, Chloroform-*d*) δ 146.85, 137.66, 134.07, 132.49 (d, *J* = 7.5 Hz), 128.97 (d, *J* = 214.3 Hz), 128.92, 128.62, 125.02, 64.34 (d, *J* = 4.3 Hz).
$^{31}$**P NMR** (243 MHz, Chloroform-*d*) δ 18.60.
**HRMS** for $C_{16}H_{16}N_2O_7P^+$ $[M+H]^+$ calc.: 379.0690 Da; found: 379.0717 Da

2-nitrobenzyl hydrogen vinylphosphonate (**5b**):

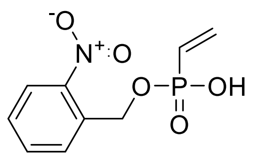

**5a** (1 mmol) was dissolved in 3 ml THF followed by the addition of 1 ml $H_2O$ and 1 ml of a 1M aq. NaOH solution. The reaction was stirred at room temperature for 2 h. After completion, the reaction mixture was diluted with $H_2O$ and extracted twice with $Et_2O$ (20 ml each). The aqueous layer was acidified with 2 M HCl and extracted with EtOAc (2x 20 ml). Combined EtOAc extracts were dried over $MgSO_4$, filtered and evaporated to obtain **5b** as an off-white solid (161 mg, 70% yield).

**$^1$H NMR** (300 MHz, DMSO-$d_6$) δ 8.14 (dd, *J* = 8.2, 1.3 Hz, 1H), 7.92 – 7.73 (m, 1H), 7.63 (ddd, *J* = 8.7, 7.2, 1.8 Hz, 1H), 6.40 – 5.89 (m, 2H), 5.29 (d, *J* = 7.6 Hz, 1H).
**$^{13}$C NMR** (75 MHz, DMSO-$d_6$) δ 147.19, 134.72, 134.19, 133.58 (d, *J* = 7.5 Hz), 129.96, 129.21 (d, *J* = 34.2 Hz), 127.60, 125.25, 63.15 (d, *J* = 3.8 Hz).
**$^{31}$P NMR** (122 MHz, DMSO-$d_6$) δ 14.83.
**HRMS** for $C_9H_{11}NO_5P^+$ $[M+H]^+$ calc.: 244.0369 Da; found: 244.0329 Da

2-nitrobenzyl N-(N-(3-phenylpropyl)carbamimidoyl)-P-vinylphosphonamidate (**5d**):

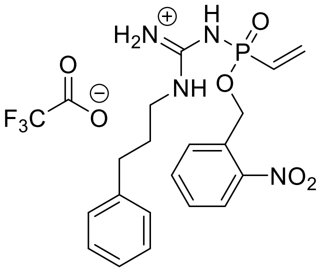

**5b** was dissolved in dry DCM (0.1 M) with a catalytic amount of DMF. 4 eq. oxalylchloride were added dropwise followed by vigorous stirring at room-temperature and occasional heating until the phosphorus NMR indicated full conversion to the corresponding chloride ($^{31}$P-NMR: δ 27.52 ppm). All volatiles were removed under argon, and the intermediate was dissolved in dry THF (0.1 M). In parallel compound **1** was dissolved in dry THF and deprotonated using 2.5 eq. Barton's base. The chloride intermediate was carefully added to deprotonated **1** and stirred for 30 min. at room temperature. 12.4 mg **5d** were obtained after purification via semi preparative HPLC (57% yield).

**$^1$H NMR** (600 MHz, DMSO-$d_6$) δ 8.42 – 8.29 (m, 1H), 8.17 (dd, *J* = 8.2, 3.4 Hz, 1H), 7.83 (dt, *J* = 11.6, 4.1 Hz, 2H), 7.66 (ddq, *J* = 8.6, 6.4, 1.9 Hz, 1H), 7.30 (td, *J* = 7.7, 3.6 Hz, 2H), 7.27 – 7.14 (m, 3H), 6.63 – 6.27 (m, 3H), 5.63 – 5.48 (m, 2H), 3.24 (qd, *J* = 6.9, 4.0 Hz, 2H), 2.67 – 2.57 (m, 2H), 1.92 – 1.75 (m, 2H).
**$^{13}$C NMR** (151 MHz, DMSO-$d_6$) δ 159.32 (q, *J* = 34.9 Hz; TFA), 155.46, 147.26, 141.46, 138.08, 134.74, 132.03 – 131.83 (m), 129.96, 129.35, 128.84, 128.71, 126.58 (d, *J* = 170.7 Hz), 126.40, 125.37, 116.53 (q, *J* = 293.8 Hz; TFA), 64.05, 41.48, 32.44, 30.05.
**$^{31}$P NMR** (243 MHz, DMSO-$d_6$) δ 17.14.
**HRMS** for $C_{38}H_{47}N_8O_8P_2^+$ $[2M+H]^+$ calc.: 805.2987 Da; found: 805.3033 Da

N-(N-(3-phenylpropyl)carbamimidoyl)-P-vinylphosphonamidic acid (**P2a**):

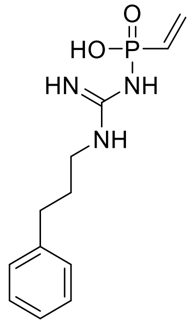

10 mg **5d** were dissolved in 2 ml degassed MeCN under argon and irradiated at 365 nm for 30 minutes with vigorous stirring (The light source used for experiments was an EvoluChem 365 nm LED (18 W, $\lambda_{max}$ = 365 nm, Hepatochem, US) in combination with the EvoluChem PhotoRedOx Box™). 1.9 mg **P2a** were obtained after purification via semi preparative HPLC (37% yield).

**$^{1}$H NMR** (600 MHz, DMSO-$d_6$) δ 8.41 (s, 1H), 7.80 (d, *J* = 68.3 Hz, 3H), 7.30 (t, *J* = 7.6 Hz, 2H), 7.21 (d, *J* = 7.5 Hz, 3H), 6.12 (td, *J* = 19.4, 12.3 Hz, 1H), 5.94 – 5.61 (m, 2H), 3.15 (q, *J* = 6.5 Hz, 2H), 2.61 (t, *J* = 7.8 Hz, 2H), 1.77 (p, *J* = 7.2 Hz, 2H).
**$^{13}$C NMR** (151 MHz, DMSO-$d_6$) δ 156.43, 141.46, 136.16 (d, *J* = 162.4 Hz), 128.86, 128.71, 127.99, 126.41, 40.55, 32.37, 30.58.
**$^{31}$P NMR** (243 MHz, DMSO-$d_6$) δ 2.97.
**HRMS** for $C_{12}H_{19}N_3O_2P^+$ $[M+H]^+$ calc.: 268.1209 Da; found: 268.1203 Da

#### Synthetic attempts towards P2b:

Initial attempts involved direct activation as well as a two-step procedure involving hydrolysis followed by activation (analogously to **P1b**). However, the direct activation with oxalylchloride was unsuccessful. In contrast, the selective mono-hydrolysis was achieved using aq. NaOH in THF. Nonetheless, despite the hydrophobic *o*-nitro-benzyl substituent, the phosphonic acid could not be transferred to the organic layer. Therefore, we changed the strategy and employed a diethylamine-derived phosphonamidate where hydrolysis with formic acid results in a volatile salt byproduct that can be removed by lyophilization from a diluted solution of triethyammonium formate. Unfortunately, after successful isolation of the phosphonic acid (**6b**) we were unable to activate it using oxalylchloride which prompted us to design a more reliable synthetic route.

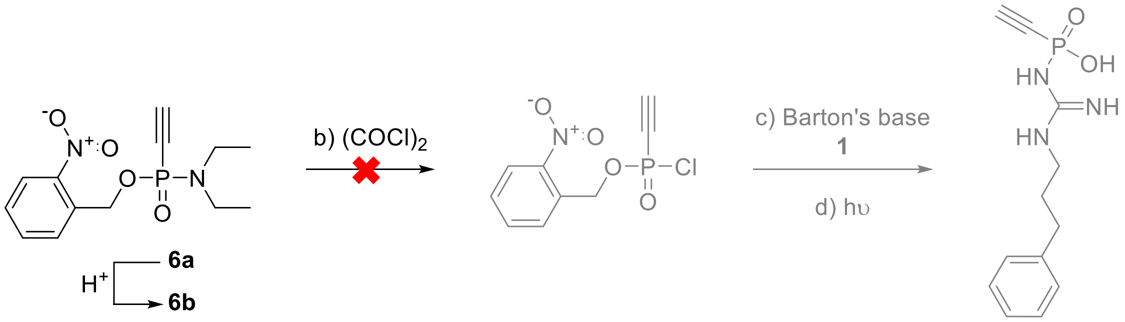

*Scheme S6: Initially proposed synthetic route towards probe **P2b***

##### 2-nitrobenzyl N,N-diethyl-P-ethynylphosphonamidate (**6a**):

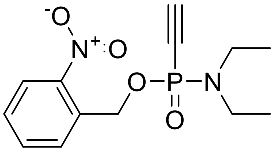

A 250-ml Schlenk flask was charged with 1750 µl $PCl_3$ (20 mmol, 1 eq.) in 100 ml of dry $Et_2O$ under an argon atmosphere, cooled to -78°C and 4.15 ml diethylamine was added drop wise. The solution was allowed to warm to room temperature and 50 ml dry hexane was added. The precipitate was filtered over cellite and the filtrate was concentrated under reduced pressure. The residue was dissolved in 20 ml dry THF and cooled to -78°C. To this solution 44 ml of ethynylmagnesium bromide (0.5 M in THF, 1.1 eq.) were slowly added and the reaction was allowed to warm to room temperature. After another 30 minutes of stirring at room temperature, it was again cooled to -78°C and 3.06 g of 2-nitrobenzyl alcohol (20 mmol, 1 eq.) dissolved in 20 ml acetonitrile and 44 ml of tetrazole dissolved in acetonitrile (0.45 M, 1 eq) were added dropwise. The mixture was allowed to warm to room temperature and was stirred overnight. The solvent was removed under reduced pressure, and the residue was dissolved in 50 ml of water extracted 3x with EtOAc. Combined organic layers were dried over $MgSO_4$, filtrated, evaporated and purified via column chromatography on silica gel (Hexane/EtOAc; 1:1) to obtain 2.526 g of **6a** as a yellow oil. (43% yield)

**$^1$H NMR (300 MHz, Chloroform-*d*)** δ 8.17 (dd, *J* = 8.2, 1.3 Hz, 1H), 7.84 (dq, *J* = 7.9, 1.1 Hz, 1H), 7.72 (td, *J* = 7.6, 1.4 Hz, 1H), 7.52 (dddd, *J* = 8.2, 7.4, 1.5, 0.7 Hz, 1H), 5.51 (qd, *J* = 15.1, 7.9 Hz, 2H), 3.23 (dqd, *J* = 12.5, 7.1, 1.1 Hz, 4H), 2.96 (d, *J* = 12.4 Hz, 1H), 1.20 (t, *J* = 7.1 Hz, 6H).
**$^{13}$C NMR (75 MHz, Chloroform-*d*)** δ 146.81, 134.09, 132.84 (d, *J* = 9.0 Hz), 128.69, 128.61, 124.98, 87.73 (d, *J* = 45.9 Hz), 76.54 (d, *J* = 261.2 Hz), 63.44 (d, *J* = 3.4 Hz), 38.93 (d, *J* = 5.6 Hz), 13.91 (d, *J* = 2.3 Hz).
**$^{31}$P NMR (122 MHz, Chloroform-*d*)** δ -0.21.
**HRMS** for $C_{13}H_{17}N_2O_4PNa^+$ [M+Na$^+$] calc.: 319.0818 Da; found: 319.0823 Da

#### 2-nitrobenzyl ethynylphosphonate triethylammonium salt (**6b**):

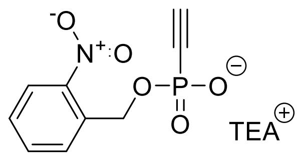

Compound **6b** was synthesized from 296 mg **6a** (1 mmol) that was hydrolyzed for 1 h in formic acid/water (95:5). After 1 h UPLC-MS indicated complete conversion and the formic acid was removed under a stream of nitrogen. The residue was lyophilized from 10 ml aqueous triethylammonium formate (100 mM, 50 vol% acetonitrile) to yield compound **6b** as a light brown solid.

**$^{1}$H NMR** (600 MHz, DMSO-$d_6$) δ 8.08 (d, *J* = 8.1 Hz, 1H), 7.81 (dt, *J* = 15.0, 7.8 Hz, 2H), 7.57 (t, *J* = 7.8 Hz, 1H), 5.17 (d, *J* = 7.7 Hz, 2H), 3.57 (d, *J* = 11.3 Hz, 1H), 2.94 (m, 6H), 1.19 (q, *J* = 7.4 Hz, 9H).
**$^{13}$C NMR** (151 MHz, DMSO-$d_6$) δ 164.63, 147.06, 134.45, 129.00, 128.85, 124.91, 84.38 (d, *J* = 41.2 Hz), 83.14 (d, *J* = 242.5 Hz), 63.16 (d, *J* = 4.0 Hz), 45.84, 8.91.
**$^{31}$P NMR** (243 MHz, DMSO-$d_6$) δ -13.54.
**HRMS** for $C_9H_8NO_5PNa^+$ $[M+Na]^+$ calc.: 264.0032 Da; found: 264.0027 Da

Based on all our previous observations we designed compound **7** that employs a *p*-nitro-phenyl leaving group instead of the rather instable and unreliable chloride. Moreover, we incorporated a triisopropyl-silyl protecting group on the terminal alkyne, to avoid potential unwanted addition of *p*-nitro-phenol after coupling to guanidinium compounds.

#### 4-nitrophenyl (1-(2-nitrophenyl)ethyl) ((triisopropylsilyl)ethynyl)phosphonate (**7**):

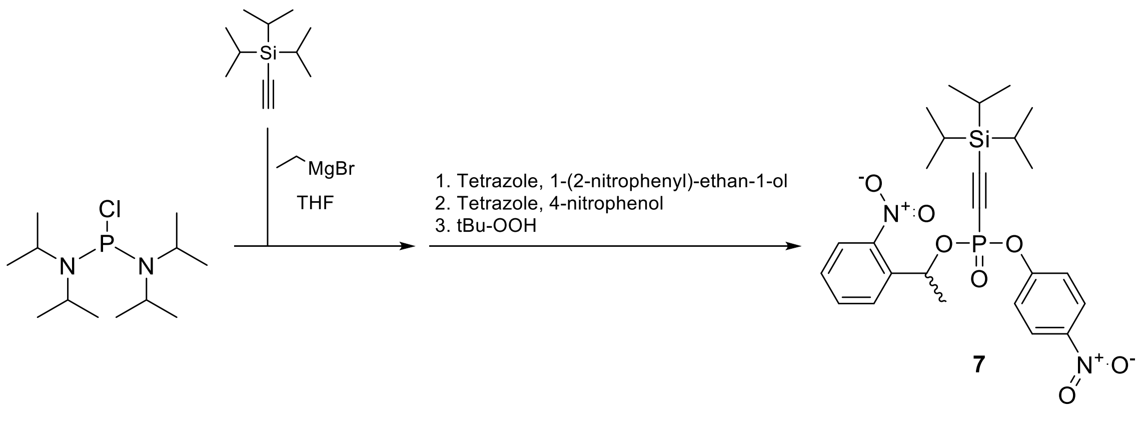

***Scheme S7**: Synthetic route towards the highly functionalized precursor **7** used for the synthesis of pArg probes.*

1.35 ml TIPS-acetylene was dissolved in 5 ml dry THF under argon and cooled on ice. Subsequently, 5 ml ethyl-magnesium bromide (1 M in THF) was added dropwise and the reaction was stirred until gas evolution has stopped. In parallel, a 100-ml round-bottom flask was charged with 1335 mg bis-(diisopropylamino)chlorophosphine (5.00 mmol) under an argon atmosphere and cooled to 0 °C. The freshly prepared TIPS-ethynyl-magnesium bromide solution (0.5 M in THF, 5.50 mmol) was added drop wise and the reaction was allowed to warm to room temperature and stirred for another 30 minutes after which the mixture was cooled to 0 °C. 1-(2-nitrophenyl)ethan-1-ol (835 mg, 5 mmol) was dissolved in 12 ml 1H-tetrazole solution (0.46 M in MeCN, 5.5 mmol, 1.1 eq.) and was added to the cooled reaction mixture. The reaction mixture was stirred 1 h at room temperature and then again cooled to 0 °C. 4-nitro phenol (695 mg, 5 mmol) was dissolved in 12 ml 1H-tetrazole solution (0.46 M in MeCN, 5.5 mmol, 1.1 eq.) and was added to the cooled reaction mixture. The reaction was allowed to warm to room temperature and stirred over night. The next day the reaction was cooled to 0 °C and 2 eq. Luperox were added slowly (1.5 ml 70% in water). Finally, the reaction-mixture was participated between 1 M aq. HCl and EtOAc. The organic layer was dried over $MgSO_4$, filtered, evaporated and purified via column chromatography on silica gel (Hexane/EtOAc; 4:1 → 2:1). The desired compound **7** was isolated as a mixture of diastereomers (360 mg, 13% yield). The reaction was repeated at a 1 mmol scale, where an improved yield of 35% could be achieved.

**$^{1}$H NMR (600 MHz, Chloroform-*d*)** δ 8.27 – 8.21 (m, 1H), 8.23 – 8.17 (m, 1H), 8.06 (dd, *J* = 8.2, 1.3 Hz, 0H), 8.03 (dd, *J* = 8.2, 1.3 Hz, 1H), 7.89 (dd, *J* = 7.9, 1.3 Hz, 0H), 7.87 (dd, *J* = 7.9, 1.3 Hz, 1H), 7.71 (dtd, *J* = 16.7, 7.7, 1.3 Hz, 1H), 7.51 (dddd, *J* = 8.6, 7.2, 5.4, 1.5 Hz, 1H), 7.44 – 7.38 (m, 1H), 7.33 – 7.28 (m, 1H), 6.43 (ddq, *J* = 31.7, 9.9, 6.3 Hz, 1H), 1.83 (d, *J* = 6.3 Hz, 3H), 1.12 – 0.97 (m, 23H).
**$^{13}$C NMR (151 MHz, Chloroform-*d*)** δ 154.50 (d, *J* = 6.4 Hz), 154.41 (d, *J* = 6.3 Hz), 146.53 (d, *J* = 41.5 Hz), 146.33, 145.02, 144.95, 136.88 (d, *J* = 4.8 Hz), 136.66 (d, *J* = 4.5 Hz), 134.17, 134.04, 129.11, 129.07, 127.84, 127.74, 125.53 (d, *J* = 7.3 Hz), 124.69, 124.52, 121.24, 121.27 – 120.93 (m), 111.57 (d, *J* = 39.2 Hz), 111.50 (d, *J* = 38.9 Hz), 93.79 (d, *J* = 283.7 Hz), 93.07, 73.71 (d, *J* = 4.8 Hz), 73.48 (d, *J* = 4.7 Hz), 24.41 (t, *J* = 5.9 Hz), 18.18 (d, *J* = 3.8 Hz), 10.64.
**$^{31}$P NMR (243 MHz, Chloroform-*d*)** δ -14.94, -15.21.
**HRMS** for $C_{25}H_{33}N_2O_7PSiNa^+$ $[M+Na]^+$ calc.: 555.1687 Da; found: 555.1692 Da

P-ethynyl-N-(N-(3-phenylpropyl)carbamimidoyl)phosphonamidic acid (**P2b**):

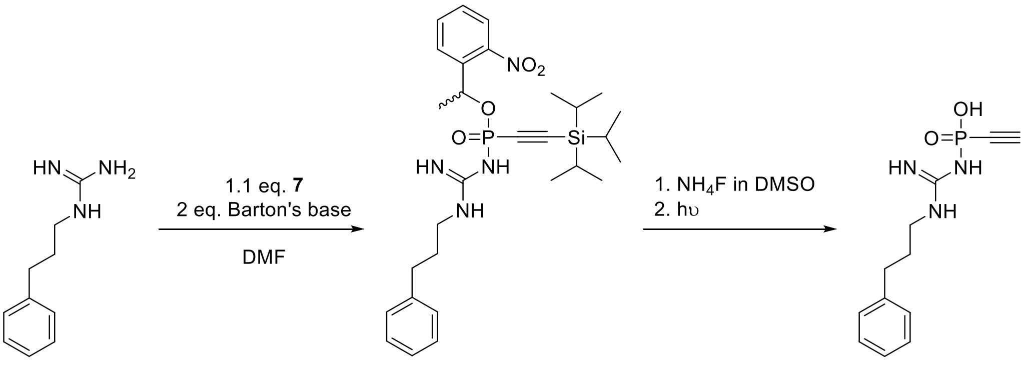

***Scheme S8**: Synthetic route towards covalent pArg-phosphatase probe **P2b**.*

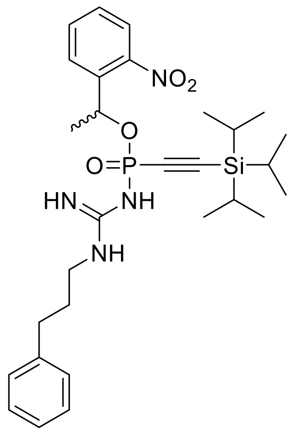

10 mg **1** were dissolved in DMF (0.15 M) alongside 2 eq. Barton's base, followed by the addition of 1.1 eq. **7** (0.1 M in DMF). After 5 minutes UPLC-MS analysis indicated full conversion and the reaction was purified via semi preparative HPLC and the obtained intermediate **8** was (12 mg, 62%) directly used in the next step.

**$^{1}$H NMR** (600 MHz, DMSO-$d_6$) δ 8.16 – 7.93 (m, 1H), 7.87 – 7.74 (m, 2H), 7.61 – 7.50 (m, 1H), 7.27 (td, *J* = 7.6, 1.9 Hz, 2H), 7.23 – 7.13 (m, 3H), 2.60 – 2.55 (m, 1H), 1.57 (dd, *J* = 8.8, 6.3 Hz, 3H), 1.07 – 0.82 (m, 23H).
**$^{31}$P NMR** (243 MHz, DMSO-$d_6$) δ -6.70, -7.24.

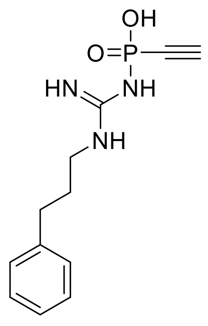

The intermediate from the first step in 0.33 ml DMSO-$d_6$ (approx. 8 mg) was diluted with MeCN & 60 µl aq. $NH_4F$ (2 M in $H_2O$, 8.5 eq.) was added and stirred 5 minutes at room temperature. Subsequently, the reaction was irradiated with a Hg (Xe) arc lamp (LOT-QuantumDesign GmbH) using a 297 nm filter with 15% transmission (Andover Inc.) for 15 minutes. Finally, the product **P2b** was purified via semi preparative HPLC (1.7 mg, 45% yield).

**$^{1}$H NMR** (600 MHz, DMSO-$d_6$) δ 8.28 (s, 1H), 8.02 (s, 1H), 7.77 (s, 2H), 7.30 (t, *J* = 7.6 Hz, 2H), 7.25 – 7.15 (m, 4H), 3.58 (d, *J* = 11.1 Hz, 1H), 3.17 (q, *J* = 6.5 Hz, 2H), 2.63 (t, *J* = 7.8 Hz, 2H), 1.79 (p, *J* = 7.6 Hz, 2H).
**$^{31}$P NMR** (243 MHz, DMSO-$d_6$) δ 16.60.
**$^{13}$C NMR** (151 MHz, DMSO-$d_6$) δ 156.02, 141.43, 128.86, 128.73, 126.40, 85.21 (d, *J* = 231.4 Hz), 83.92 (d, *J* = 40.1 Hz), 40.56, 32.31, 30.47.
**HRMS** for $C_{12}H1_7N_3O_2P$ $[M+H]^+$ calc.: 266.1053 Da; found: 266.1075 Da

### Synthesis of functional pArg-phosphatase probes:

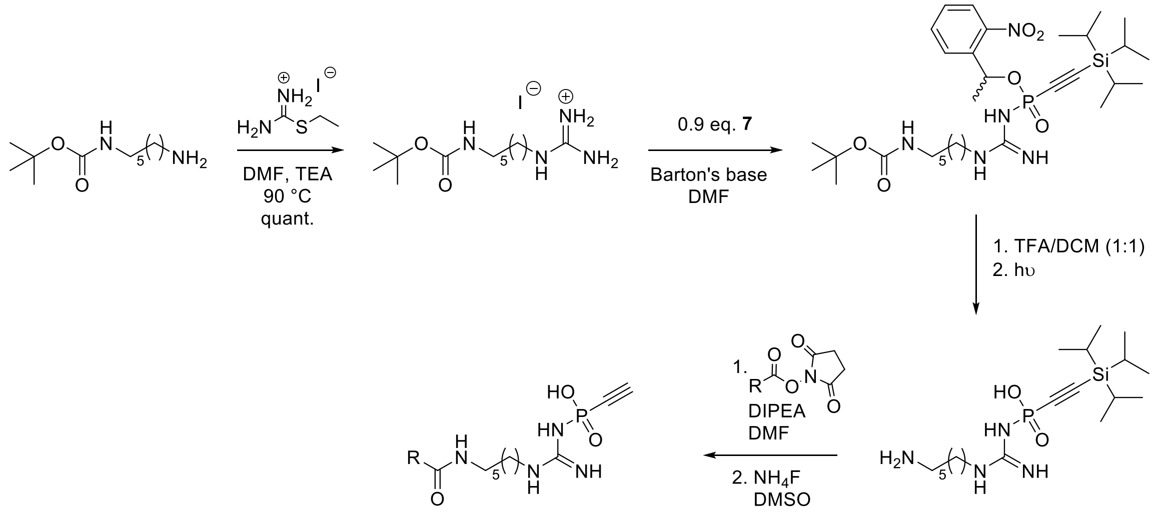

***Scheme S9**: Synthetic route towards functionalized pArg-phosphatase probes*

amino((6-((tert-butoxycarbonyl)amino)hexyl)amino)methaniminium iodide (**9**):

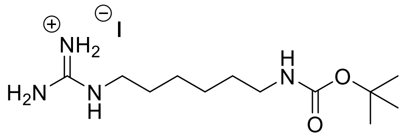

1080 mg of *N*-Boc-1,6-hexanediamine (5 mmol) were dissolved in 1 ml DMF and 1160 mg S-ethyl isothiourea hydroiodide (5 mmol) were added. The reaction was stirred at 90 °C over night (open to air). After removal of the solvent, **9** was obtained as a clear viscous liquid in quantitative yield.

**$^1$H NMR** (600 MHz, DMSO-$d_6$) δ 6.72 (t, *J* = 5.7 Hz, 1H), 3.08 (t, *J* = 7.1 Hz, 2H), 2.94 – 2.85 (m, 2H), 1.43 (p, *J* = 7.2 Hz, 2H), 1.35 (s, 11H), 1.25 (qd, *J* = 12.2, 11.3, 7.2 Hz, 4H).
**$^{13}$C NMR** (151 MHz, DMSO-$d_6$) δ 157.06, 156.02, 77.77, 41.20, 40.19, 29.82, 28.79 (d, *J* = 15.1 Hz), 26.31, 26.18.
**HRMS** for $C_{12}H_{27}N_4O_2$ $[M+H]^+$ calc.: 259.2129 Da; found: 259.2115 Da

tert-butyl (8-imino-3,3-diisopropyl-2-methyl-6-(1-(2-nitrophenyl)ethoxy)-6-oxo-7,9-diaza-6l5-phospha-3-silapentadec-4-yn-15-yl)carbamate (**10**):

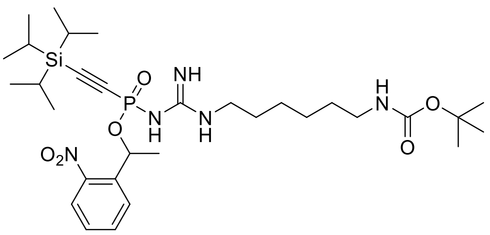

To a stirring solution of 4-nitrophenyl (1-(2-nitrophenyl)ethyl) ((triisopropylsilyl)ethynyl)phosphonate (**7**, 109 mg, 0.2 mmol, 1 eq) in DMSO (0.5 mL) was added, a separately prepared solution of amino((6-((tert-butoxycarbonyl)amino) hexyl)amino)methaniminium iodide (**9**, 77 mq, 0.2 mmol, 1 eq), and Barton's Base (51 mg, 0.3 mmol, 1.5 eq) in 0.5 mL DMSO. The resultant mixture was stirred at room temperature and monitored via UPLC-MS for consumption of 4-

nitrophenyl (1-(2-nitrophenyl)ethyl) ((triisopropylsilyl)ethynyl)phosphonate, where after complete consumption the mixture was diluted and purified directly via prep-HPLC (10 – 95% MeCN in $H_2O$/0.1% TFA over 65 min) to afford the title compound as a brown solid composed of a mixture of diastereomers in a 72% yield.

**$^{1}$H NMR** (600 MHz, DMSO-$d_6$) δ 8.54 – 8.32 (m, 6H), 8.02 – 7.96 (m, 2H), 7.94 – 7.89 (m, 1H), 7.78 (dtd, *J* = 29.2, 7.7, 1.4 Hz, 3H), 7.71 (dtd, *J* = 12.7, 7.6, 1.3 Hz, 2H), 7.62 (ddd, *J* = 8.5, 7.3, 1.5 Hz, 1H), 7.60 – 7.53 (m, 4H), 7.20 (dd, *J* = 17.9, 15.0 Hz, 1H), 6.80 – 6.73 (m, 2H), 5.95 – 5.89 (m, 1H), 5.81 (dq, *J* = 8.9, 6.3 Hz, 1H), 5.74 (dq, *J* = 8.7, 6.3 Hz, 1H), 5.23 (dd, *J* = 15.0, 11.0 Hz, 1H), 3.61 (t, *J* = 7.7 Hz, 1H), 3.49 (dddd, *J* = 35.8, 15.4, 9.5, 5.9 Hz, 2H), 2.90 (hept, *J* = 5.5 Hz, 4H), 1.61 (dd, *J* = 14.6, 6.3 Hz, 4H), 1.51 – 1.45 (m, 8H), 1.40 – 1.30 (m, 21H), 1.28 – 1.22 (m, 3H), 1.18 (d, *J* = 7.1 Hz, 4H).
**$^{13}$C NMR** (151 MHz, DMSO-$d_6$) δ 159.17, 158.95, 158.73, 158.51, 157.24, 157.08, 156.90, 156.06, 147.29, 147.07, 147.00, 146.95, 146.80, 143.37, 143.22, 141.34, 137.45, 137.43, 137.40, 137.26, 137.23, 137.18, 134.55, 134.53, 134.48, 134.30, 129.75, 129.72, 129.63, 129.60, 129.32, 129.00, 128.82, 128.39, 128.32, 128.06, 127.97, 127.92, 124.79, 124.76, 124.67, 124.25, 118.02, 116.05, 96.31, 95.88, 94.99, 94.57, 77.79, 70.06, 69.94, 69.91, 69.88, 68.04, 45.86, 45.74, 45.64, 40.55, 40.42, 40.28, 40.15, 40.01, 39.87, 39.73, 39.59, 29.91, 28.90, 28.73, 26.52, 26.46, 26.32, 26.14, 25.99, 25.81, 25.77, 25.03, 24.98, 24.69, 24.66, 24.61, 24.57, 24.52, 24.49, 18.31, 12.55.
**$^{31}$P NMR** (243 MHz, DMSO-$d_6$) δ 20.20, 20.03, 19.98, 8.27, 7.70.
**HRMS** for $C_{31}H_{54}N_5O_5PSi^+$ $[M+H]^+$ calc.: 652.3654 Da; found: 663.2884 Da $[2M+Na]^{2+}$

N-(N-(6-aminohexyl)carbamimidoyl)-P-((triisopropylsilyl)ethynyl)phosphonamidic acid (**11**):

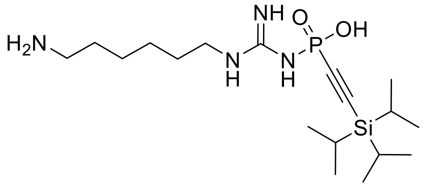

**10** was dissolved in a mixture of TFA/DCM (1:1) and stirred for 30 minutes at room temperature. Subsequently, all volatiles were removed under a stream of argon, followed by lyophilization from water/MeCN (1:1). The crude intermediate was dissolved in 2 ml degassed MeCN under argon and irradiated at 365 nm for 30 minutes with vigorous stirring (The light source used for experiments was an EvoluChem 365 nm LED (18 W, $\lambda_{max}$ = 365 nm, Hepatochem, US) in combination with the EvoluChem PhotoRedOx Box™). The obtained intermediate **10c** was used in the next step without further purification. The crude reaction mixture containing intermediate **10c** was aliquoted for subsequent reaction with functional NHS-esters. Additionally, a small portion of the reaction mixture was purified for analytical purposes.

**$^{1}$H NMR** (600 MHz, DMSO-$d_6$) δ 8.05 (s, 1H), 7.69 (s, 4H), 3.16 (q, *J* = 6.6 Hz, 2H), 2.85 – 2.72 (m, 2H), 1.50 (dt, *J* = 36.4, 7.1 Hz, 4H), 1.31 (p, *J* = 3.6 Hz, 4H), 1.04 (d, *J* = 2.4 Hz, 21H).
**$^{31}$P NMR** (243 MHz, DMSO-$d_6$) δ -23.39.
**$^{13}$C NMR** (151 MHz, DMSO-$d_6$) δ 156.06, 109.47 (d, *J* = 215.9 Hz), 94.54 (d, *J* = 30.5 Hz), 40.95, 39.19, 28.54, 27.28, 25.87, 25.77, 18.81 (9C), 10.95 (3C).
**HRMS** for $C_{18}H_{40}N_4O_2PSi^+$ $[M+H]^+$ calc.: 403.2653 Da; found: 403.2640 Da

**General procedure for functionalization & TIPS-deprotection of pArg-phosphatase probes:**

An aliquot of **10c** was mixed with 1.1 eq. of the corresponding NHS-ester dissolved in DMSO followed by the addition of 2.5 eq. DIPEA. After full consumption of the NHS-ester the reaction mixture was purified via semi preparative HPLC. The purified intermediate was dissolved in dry DMSO (at a concentration of 10 mM) and 20 eq. $NH_4F$ (2 M in $H_2O$) were added. After 30 minutes the reaction was checked by UPLC-MS. In case the reaction was not complete yet, another 2 eq. $NH_4F$ were added. Otherwise the products were purified via semi preparative HPLC.

P-ethynyl-N-(N-(6-(5-(2-oxohexahydro-1H-thieno[3,4-d]imidazol-4-yl)pentanamido)hexyl) carbamimidoyl)phosphonamidic acid **[Biotin-pArg-Phosphatase Probe, P3]:**

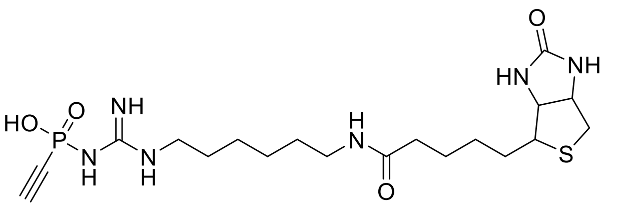

The probe was synthesized according to procedure from 5 mg Biotin-NHS and obtained in 63% yield after the second HPLC purification (4.5 mg, 40% yield from *N*-Boc-1,6-hexane diamine)

**$^1$H NMR** (600 MHz, DMSO-$d_6$) δ 4.32 (dd, *J* = 7.7, 4.9 Hz, 1H), 4.14 (dd, *J* = 7.7, 4.4 Hz, 1H), 3.57 (d, *J* = 11.3 Hz, 1H), 3.15 (q, *J* = 6.6 Hz, 2H), 3.10 (ddd, *J* = 8.8, 6.0, 4.4 Hz, 1H), 3.02 (q, *J* = 6.6 Hz, 2H), 2.83 (dd, *J* = 12.4, 5.1 Hz, 1H), 2.61 – 2.54 (m, 1H), 2.05 (t, *J* = 7.4 Hz, 2H), 1.69 – 1.19 (m, 14H).
**$^{13}$C NMR** (151 MHz, DMSO-$d_6$) δ 172.36, 163.20, 155.92, 84.99 (d, *J* = 233.0 Hz), 84.04 (d, *J* = 41.0 Hz), 61.54, 59.68, 55.91, 41.07, 38.76, 35.68, 29.49, 28.68, 28.51, 26.40, 26.10, 25.81.
**$^{31}$P NMR** (243 MHz, DMSO-$d_6$) δ -23.16.
**HRMS** for $C_{19}H_{34}N_6O_4PS^+$ $[M+H]^+$ calc.: 473.2094 Da; found: 473.2106 Da

5-((6-(3-(ethynyl(hydroxy)phosphoryl)guanidino)hexyl)carbamoyl)-2-(6-hydroxy-3-oxo-3H-xanthen-9-yl)benzoic acid **[Fluorescein-pArg-Phosphatase Probe, P4]**:

From 3.7 mg NHS-Fluorescein (3.2 mg, 44% yield from *N*-Boc-1,6-hexanediamine)

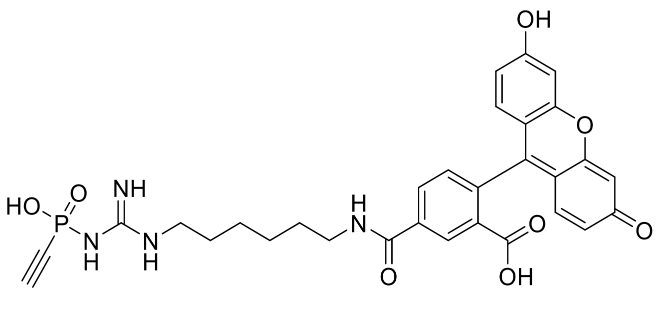

**$^{31}$P NMR** (243 MHz, DMSO-$d_6$) δ -22.68.
**HRMS** for $C_{30}H_{30}N_4O_8P^+$ $[M+H]^+$ calc.: 605.1796 Da; found: 605.1815 Da

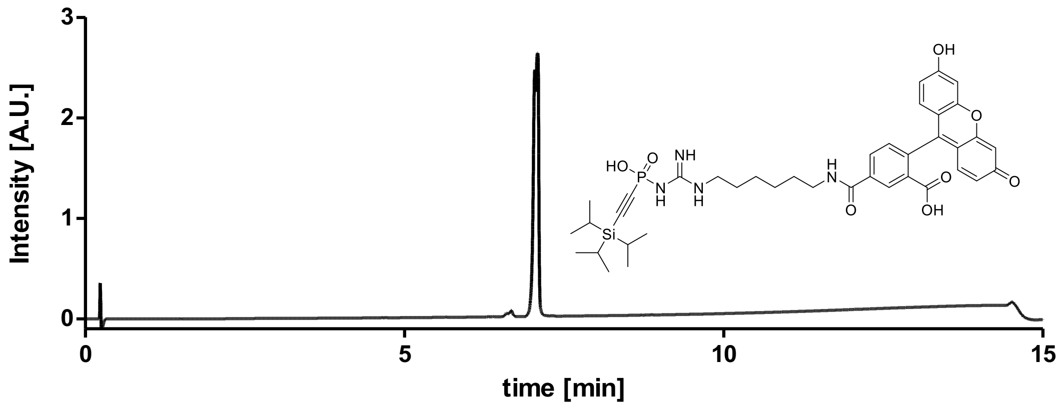

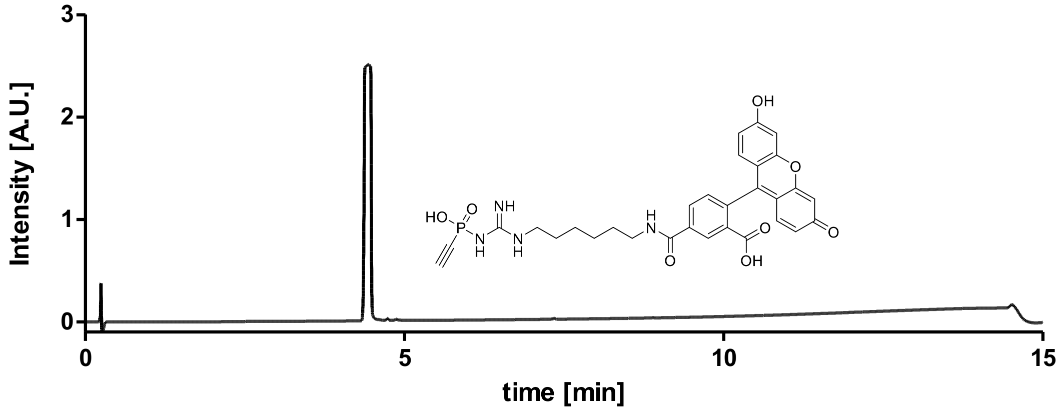

N-(9-(2-carboxy-4-((6-(3-(ethynyl(hydroxy)phosphoryl)guanidino)hexyl)carbamoyl)phenyl)-6-(dimethylamino)-3H-xanthen-3-ylidene)-N-methylmethanaminium
**[TAMRA-pArg-Phosphatase Probe, P5]:**

From 9.5 mg NHS-TAMRA 5/6-isomer (6.1 mg, 33% yield from *N*-Boc-1,6-hexanediamine)

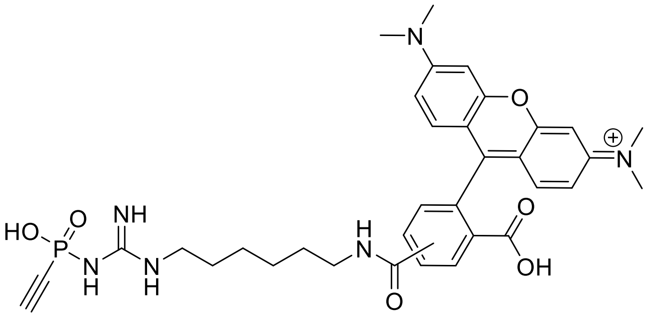

$^{31}$**P NMR** (243 MHz, DMSO-$d_6$) δ -23.09.
**HRMS** for $C_{34}H_{40}N_6O_6P^+$ $[M+H]^+$ calc.: 659.2741 Da; found: 659.2715 Da

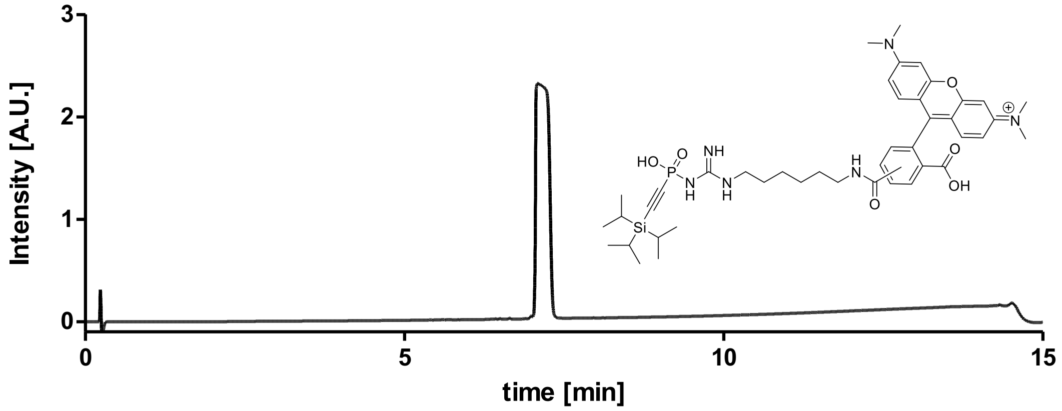

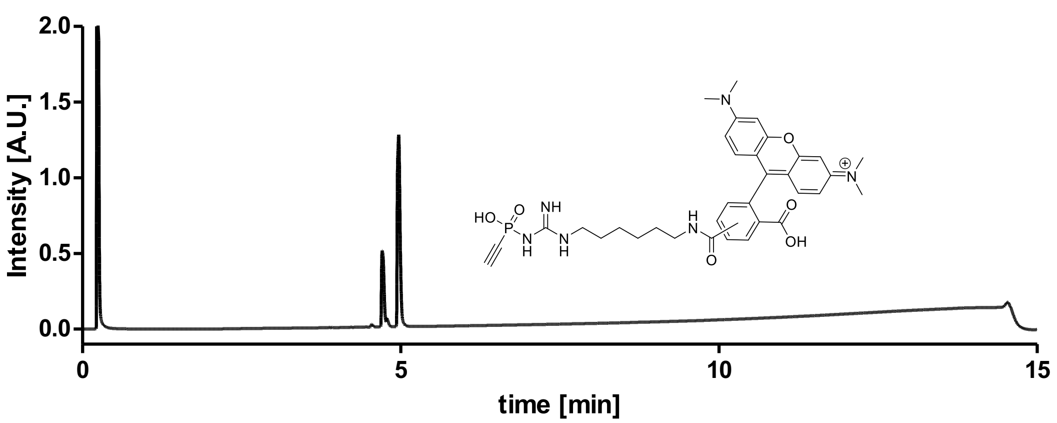

P-ethynyl-N-(N-(6-(4-(6-methyl-1,2,4,5-tetrazin-3-yl)benzamido)hexyl)carbamimidoyl) phosphonamidic acid (**P6**):

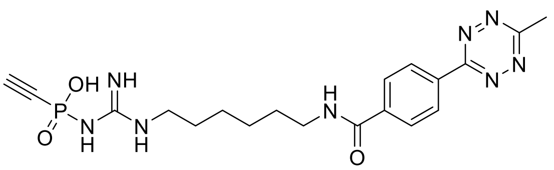

To a solution of N-(N-(6-(4-(6-methyl-1,2,4,5-tetrazin 3-yl)benzamido)hexyl)carbamimidoyl)-P-((triiso propylsilyl) ethynyl)phosphonamidic acid (10 mM, 10.4 mg) in DMSO in a 50 mL Falcon tube was added $NH_4F$ (200 mM in $H_2O$, 20 eq) and the reaction was shaken at 220 rpm for 30 min (complete consumption of starting material via UPLC-MS). The reaction mixture was directly diluted and purified via prep-HPLC (5 – 95% MeCN in $H_2O$/0.1% TFA over 55 min) to afford the title compound as a pink solid (7.6 mg, 41% yield from *N*-boc-1,6-diaminohexane).

**$^1$H NMR** (600 MHz, DMSO-$d_6$) δ 8.69 (t, *J* = 5.6 Hz, 1H), 8.54 (d, *J* = 8.0 Hz, 2H), 8.10 (d, *J* = 8.4 Hz, 2H), 8.00 (s, 1H), 7.77 (s, 1H), 3.61 (s, 2H), 3.57 (dd, *J* = 11.3, 2.9 Hz, 2H), 3.31 (q, *J* = 6.7 Hz, 2H), 3.18 (q, *J* = 6.6 Hz, 2H), 3.02 (s, 3H), 1.57 (q, *J* = 7.0 Hz, 2H), 1.50 (q, *J* = 7.2 Hz, 2H), 1.40 – 1.33 (m, 4H).
**$^{13}$C NMR** (151 MHz, DMSO-$d_6$) δ 167.79, 167.74, 165.86, 163.38, 155.95, 138.47, 138.42, 138.37, 134.54, 129.20, 129.16, 128.35, 128.31, 128.12, 128.07, 127.26, 127.22, 84.98, 84.71, 84.43, 83.35, 83.08, 40.42, 40.28, 40.14, 40.00, 39.86, 39.72, 39.58, 30.23, 29.40, 28.57, 27.86, 27.33, 26.98, 26.52, 26.16, 25.70, 25.33, 22.64, 21.78, 20.92, 20.06.
**$^{31}$P NMR** (243 MHz, DMSO-$d_6$) δ -23.08, -23.13.
**UPLC-MS** retention time (rt) = 4.05 min (>95%, 5 – 95% MeCN in $H_2O$/0.1% TFA over 15 min)
**HRMS** for $C_{19}H_{27}N_8O_3P^+$ $[M+2H]^+$ calc.: 446.1927 Da; found: 446.1898 Da

### 3. Peptide Synthesis

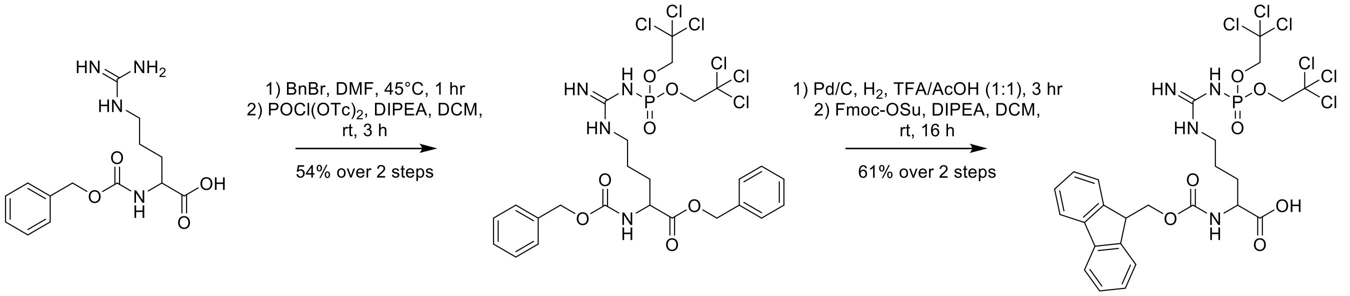

***Scheme S10**: The synthesis of Fmoc-cpArg-OH.*

**Z-Arg(PO(OTCe)$_2$)-OBn:**

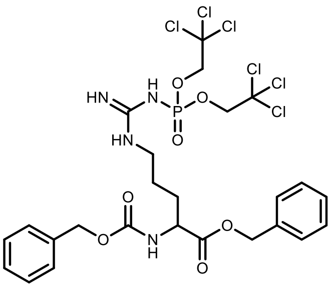

Z-Arg(PO(OTCe)$_2$)-OBn was accessed using procedures established by Seebeck and co-workers. The title product was accessed as a low melting point colourless solid in a 54% yield. All spectral data is in perfect agreement with literature.[1]

**Fmoc-Arg(PO(OTCe)$_2$)-OH:**

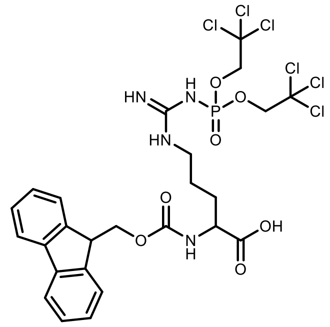

Fmoc-Arg(PO(OTCe)$_2$)-OH was accessed using procedures established by Seebeck and co-workers. The title product was accessed as a beige solid in a 61% yield. All spectral data is in perfect agreement with literature.[7]

**$NH_2$-TTAVEIDpYDSLK-OH**

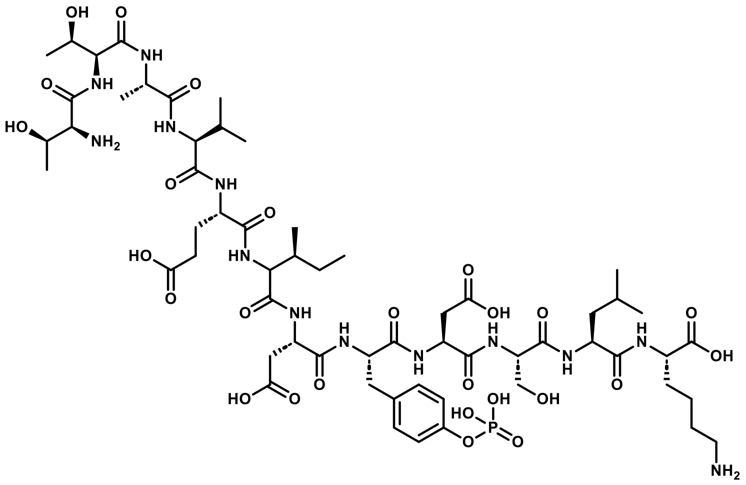

**UPLC-MS** retention time (rt) = 3.83 min (91%, 5 – 95% MeCN in $H_2O$/0.1% TFA over 15 min).
**HRMS** for $C_{59}H_{97}N_{13}O_{26}P^+$ $[M+H]^+$ calc.: 1434.6400 Da; found: 1434.6130 Da $[M+H]^+$.

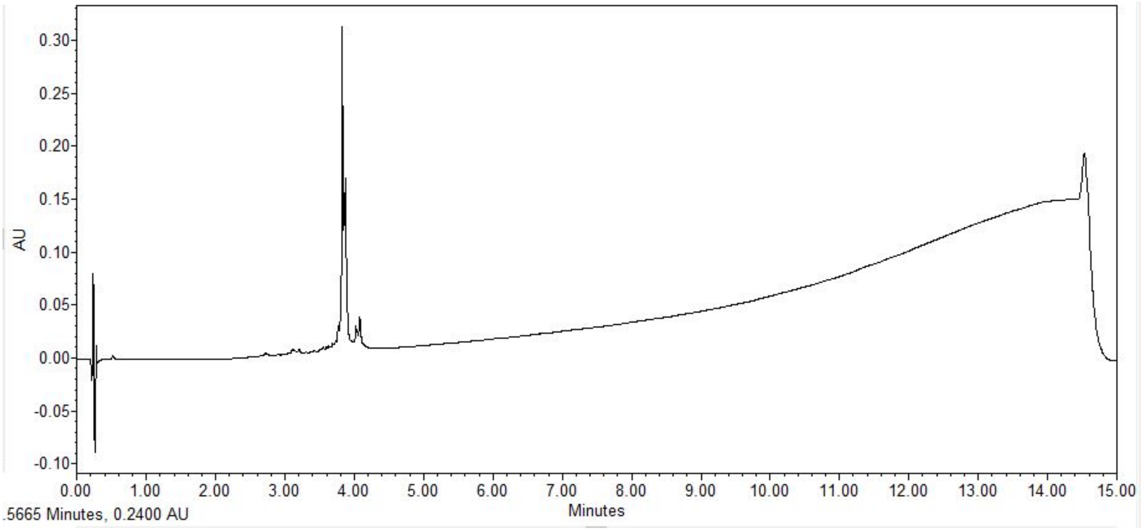

**NH$_2$-TTAVEIDcpRDSLK-OH**

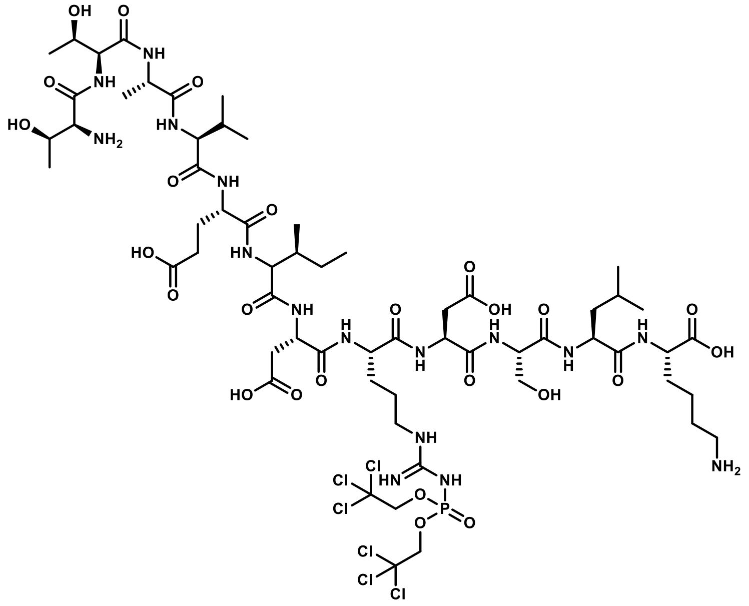

**UPLC-MS** retention time (rt) = 5.73 min (5 – 95% MeCN in $H_2O$/0.1% TFA over 15 min)
**HRMS (ESI-TOF)** for $C_{60}H_{102}N_{61}Cl_6O_{25}P^+$ $[M+H]^+$ calc.: 1687.5065 Da; found: 1690.5280 Da $[M+3H]^+$.

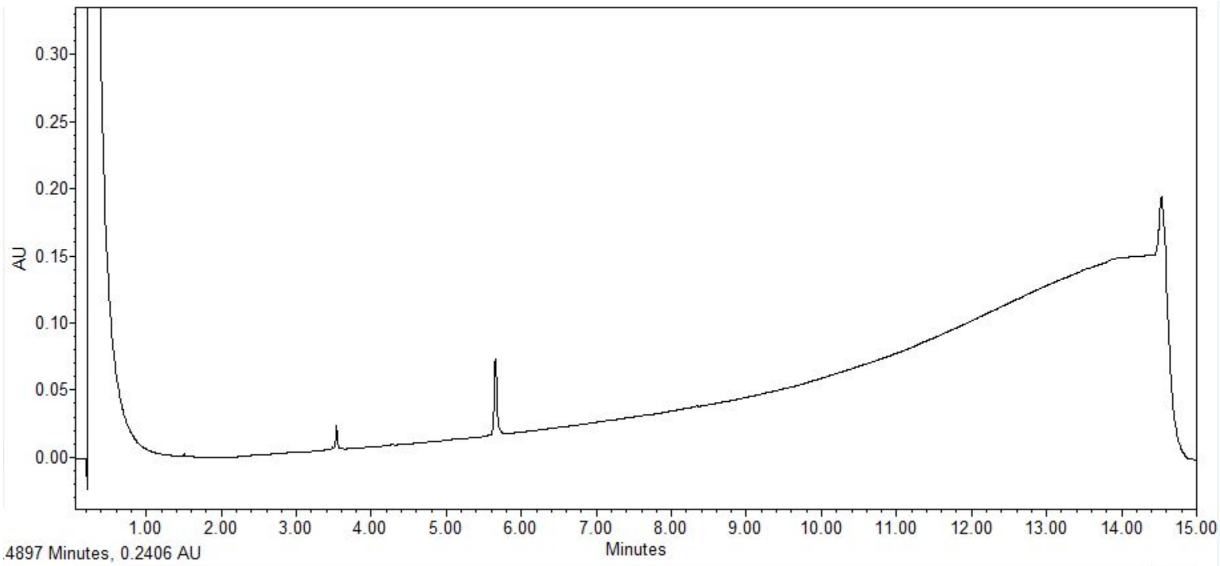

**Note:** Arg(PO(OTCE)$_2$) residue is highly acid labile and readily hydrolyses under acidic conditions present in UPLC-MS mobile phases to afford a second peak (rt – 3.534 min) corresponding to the des-phosphorylated species (17% hydrolyzed under UPLC conditions).

cpArg deprotection was carried out in lieu of experiments and kept dry at -70 °C until needed. For deprotection, NH$_2$-TTAVEIDcpRDSLK-OH (11.9 mg, 7.05 µmol) was dissolved in an 80:20 mixture of EtOH and 100 mM $NH_4CO_3$ (dd$H_2O$, pH 8.2, 3.5 mL). This mixture was sealed and subsequently subjected to a series of three freeze-pump-thaw cycles with Ar, and under a gentle stream of Ar a catalytic quantity of Pd/C (approx. 2 mg, 10-20 mol%). The system was again sealed and evacuated with Ar. Following this, whilst stirring vigorously, Ar was bubbled into the solvent for 5 min. Thereafter, the balloon was exchanged for $H_2$, which was again bubbled for 5 min. The atmosphere in the reaction vessel was exchanged for $H_2$, and this atmosphere was maintained whilst stirring overnight at room temperature. Reaction progress was monitored via UPLC-MS (5-95% MeCN in $H_2O$/0.1% TFA), and once complete the mixture was filtered through a 0.2 µM syringe to remove the catalyst. The filtrate was lyophilized to afford the deprotected peptide which was used without further purification.
